# Microtubule anchoring and coupling of CD20 to the RhoA/Rock1 pathway

**DOI:** 10.1101/2025.05.29.656301

**Authors:** Kathrin Kläsener, Cindy Eunhee Lee, Julian Bender, Angela Naumann, Lena Reimann, Geoffroy Andrieux, Claudio Mussolino, Nadja Herrmann, Roland Nitschke, Reinhard E. Voll, Bettina Warscheid, Klaus Warnatz, Michael Reth

## Abstract

CD20 is a B cell-specific four-helix transmembrane protein and a prominent target of therapeutic anti-CD20 antibodies. CD20 is localized within a membrane nanocluster harboring the IgD class B cell antigen receptor (IgD-BCR) where it functions as a gatekeeper for the resting state of naïve B lymphocytes. How CD20 exerts its gatekeeper function is not yet known. Using Ramos and human peripheral blood B cells, we show here that the serine/threonine kinase PKCδ, constitutively phosphorylates serine residues at the cytosolic tails of CD20. The phosphorylated CD20 becomes a target for 14-3-3 adaptor proteins that link CD20 to the RhoA GDP/GTP exchange factor GEF-H1 and the microtubule (MT) network controlling the stability of the IgD-BCR nanocluster. The binding of anti-CD20 antibodies results in MT disassembly and the replacement of the GEF-H1/CD20 complex by a RhoA-GTP/ROCK1/CD20 complex which drives actomyosin contractility. Our study suggests that CD20 not only maintains the resting state, but also orchestrates the MT/actin switch in active B lymphocytes. This could have implications for treatment with anti-CD20 antibodies and may help to optimize therapeutic protocols.

**Graphic abstract:** 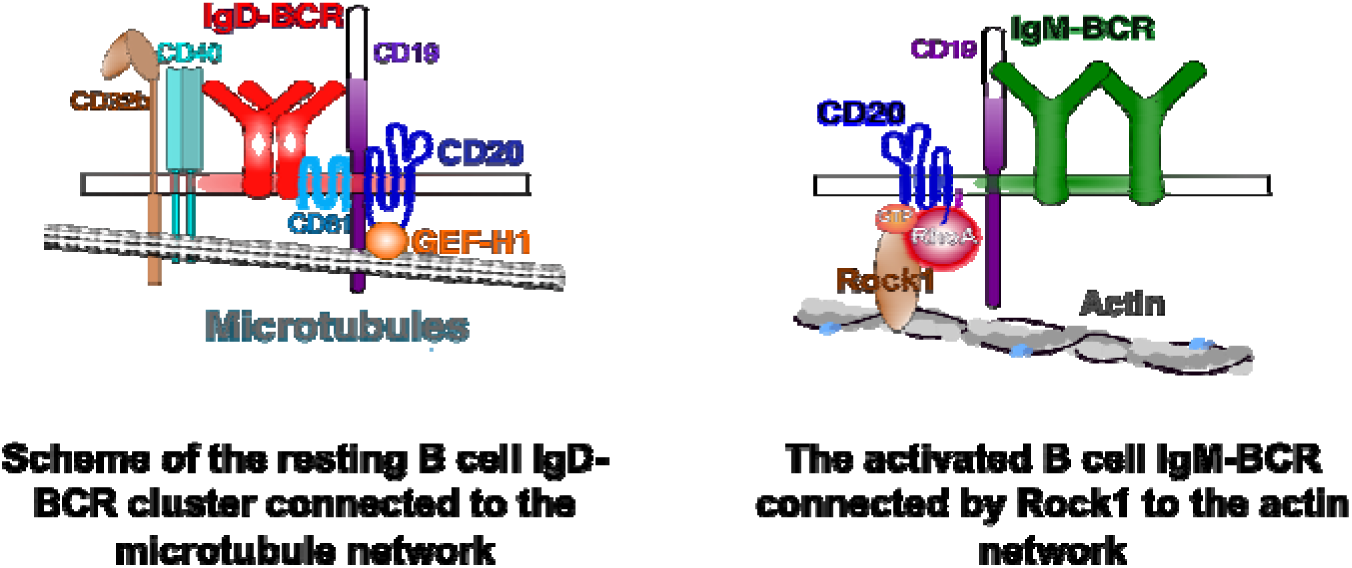

## Introduction

Upon a successful production of an immunoglobulin heavy and light chain pair, pre-B cells become immature B cells expressing an IgM-class B cell antigen receptor (IgM-BCR). The IgM-BCR plays an important role for the central tolerance process where immature B cells undergo negative selection to eliminate self-reactive clones. After surviving selection, immature B cells further develop into transitional B cells that migrate to secondary lymphoid organs and then into mature naive B cells co-expressing an IgM and IgD-BCR^1,2^. According to the clonal selection theory, mature B cells only become fully activated upon binding to their cognate antigen. Thus, most mature naive B cells remain in a dormant state and the IgD-BCR plays an important role for the maintenance of this resting state. We have previously shown that the IgM-BCR and IgD-BCR are localized in different membrane nanoclusters^3^. The underlying cytoskeleton plays an important role in the location and stability of these nanoclusters^4,5^ The IgD-BCR nanocluster has a raft-type lipid composition and harbors important B cell coreceptors such as CD19, CD20, CD40 and the BAFF receptor^6–9^ Interestingly, most proteins residing within the IgD nanocluster including the IgD-BCR reach their full expression level only at the mature naive B cell stage. This is also the case for CD20, a 37 kD non-glycosylated phosphoprotein with four transmembrane domains that form dimers or higher oligomers on the B cell surface^10–12^. The cDNA of human CD20 was first cloned in 1988^13^ and found to be encoded by the *MS4A1* gene that is one of the 18 members of the *MS4A* gene family. Due to the lack of a known ligand the biological function of CD20 has been regarded as an enigma of B cell biology^14,15^. However, newer data show that CD20 is a highly interactive protein that helps in the regulation and functional organization of other receptors on the B cell surface^16^. For example, CD70 has recently been shown to require the presence of CD20 for immune synapse formation and efficient T: B collaboration^17^.

CD20 is best known as a prominent target of therapeutic monoclonal antibodies (mAb) such as rituximab (RTX), obinutuzumab (GA101), or ocrelizumab used for the treatment of B cell chronic lymphocytic leukemia (B-CLL), malignant B cell lymphomas or human autoimmune diseases such as rheumatoid arthritis (RA), granulomatosis with polyangiitis (GPA), systemic lupus erythematosus (SLE) and, more recently, multiple sclerosis (MS)^18–23^. The anti-CD20 mAbs are thought to eliminate targeted B cells by antibody-dependent cellular cytotoxicity (ADCC) mechanisms, phagocytosis^24^, or induction of cell cycle arrest and initiation of an apoptosis program in some B cell lines^25^. The success of the anti-CD20 immunotherapy could be further improved by a more profound insight into the biological function of CD20. An important advance for this goal is the finding that CD20 is part of the IgD-BCR nanocluster where it functions as a gatekeeper for the resting state of naïve human B lymphocytes^26^. Here, we show that constitutively phosphorylated serine residues in the N- and C-terminal cytosolic tails of CD20 play an important role for this gatekeeper function by coupling CD20 to the microtubule (MT) network and the RhoA/ROCK1 signaling pathway.

## Results

### CD20 is a substrate of PKCδ, acting as a gatekeeper for the resting state of B cells

The resting state of naïve human B lymphocytes is maintained by several negative regulators including PKCdelta (PKCδ), a member of the novel serine/threonine protein kinase C (PKC) family^27,28^. PKCδ has been found to be constitutively associated with the BCR signaling subunit Igα (CD79a) and negatively regulates the BCR signaling^29^. To study the gatekeeper function of PKCδ we employed the CRISPR/Cas9 method to generate two PKCδ-deficient (PKCδ KO) clones of the human Burkitt lymphoma cell line Ramos (Fig. S1A and B). The PKCδ KO Ramos cells were viable and could be maintained in culture. We first compared the expression of B cell marker proteins on the surface of the PKCδ KO line to the wild-type (wt) Ramos cells by flow cytometric analysis (Fig. 1A). Normalization of the protein expression levels on PKCδ KO to Ramos wt cells shows that the loss of PKCδ is associated with a 25-fold increase of the B cell activation marker CD69 on Ramos B cells. The PKCδ deficiency also results in the upregulation of membrane proteins that are associated with the IgD-class nanocluster on resting B cells, namely CD20, CD19, CD40 and the IgD-BCR, whereas proteins that are associated with the IgM-BCR, such as CD45 are downregulated. In addition, two membrane proteins that function as negative B cell signaling regulators, namely CD22 and the IgG-binding Fc receptor FcγRIIb are upregulated in the PKCδ KO Ramos line (Fig. 1A). The 20-fold upregulation of the inhibitory FcγRIIb (CD32b) is interesting because mutations of CD32b are associated with human autoimmune diseases such as systemic lupus erythematosus (SLE)^30^. In summary, this study shows that PKCδ KO Ramos B cells are activated and counteract this activation by upregulating the IgD-class nanocluster and negative signaling regulators. These findings support the notion that, similar to CD20, PKCδ acts as a gatekeeper for the “resting state” of mature human B cells^27^.

**Figure 1:**
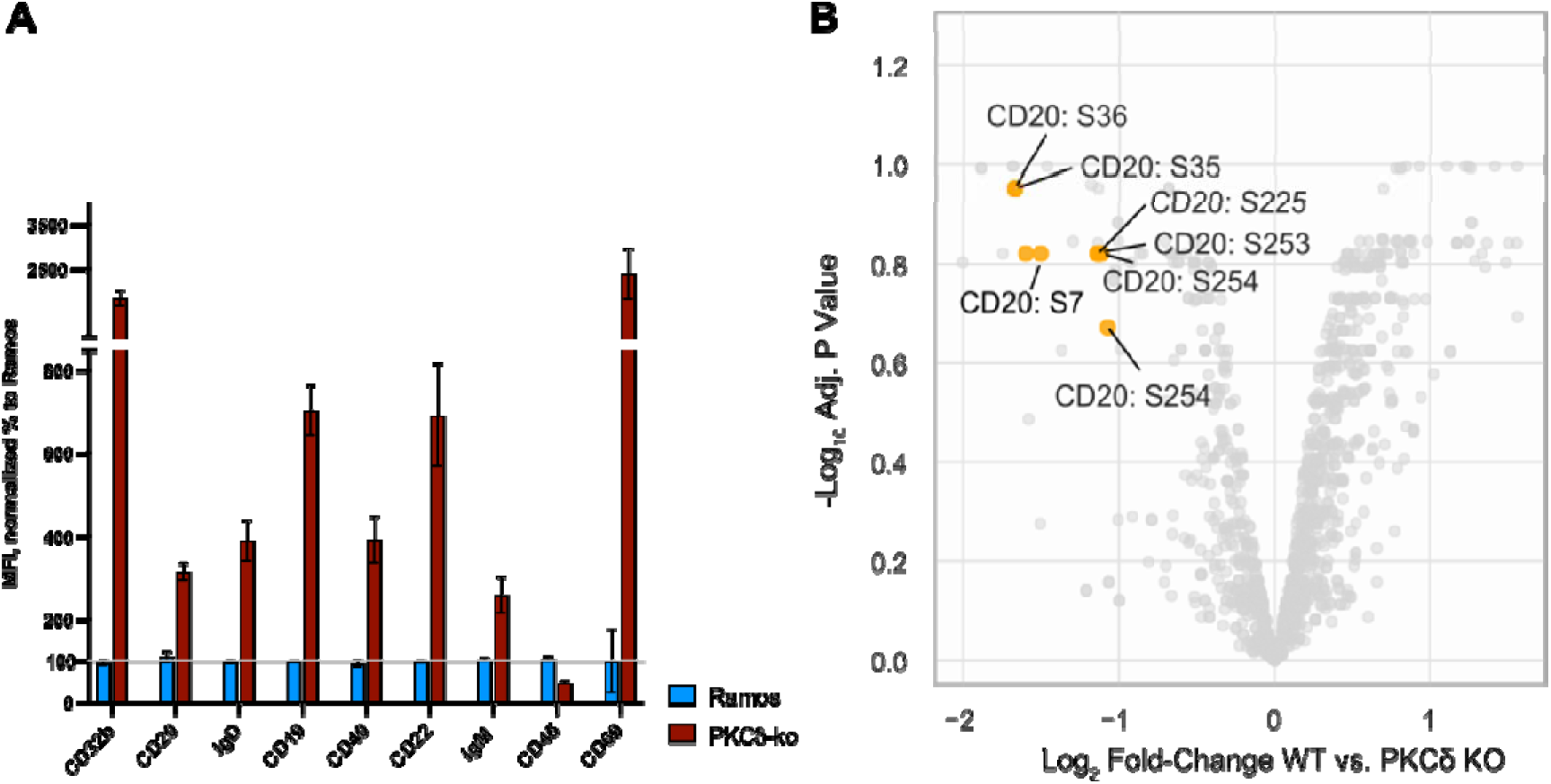
Phenotype of PKCδ KO Ramos B cells and identification of PKCδ substrate proteins. Comparison of the expression of B cell marker protein on the surface of PKCδ KO (brown) and WT Ramos B cells (blue) by flow cytometry. The expression data of PKCδ KO are normalized to WT Ramos cells, set to 100%. **B**. Volcano plot showing quantitative changes of constitutive serine/threonine phosphorylation of substrate proteins of WT in comparison to PKCδ KO Ramos B cells. Plot X-axis depicts log2 fold change of phosphorylated serine or threonine residues in the whole proteome of the two cell lines. The Y-axis shows the log10 FDR adjusted p-value <0.05. The significant changes of phosphorylated CD20 serine residues are indicated in orange.

Next, we conducted a phosphoproteome analysis to monitor the difference in the level of constitutive serine/threonine phosphorylation in Ramos wt and the PKCδ KO cell line (Fig. 1B). In this analysis we identified several serine residues in the N-terminal and C-terminal cytosolic tail of CD20 that are constitutively phosphorylated by PKCδ. Thus, PKCδ and CD20, two gatekeepers of the resting state of B lymphocytes, are connected by an enzyme/substrate relationship. Global serine/threonine phospho-proteome analysis of Ramos wt and PKCδ KO cells also reveal specific alterations in the phosphorylation of components of the RhoA/Rock-1 pathway and the MT cytoskeletal network described below (Fig. S1C). A comparison of the RNA-seq data obtained from the two cell lines indicated that, compared to Ramos wt, the PKCδ KO cells displayed an increased expression of genes encoding for proteins involved in MT depolymerization and a loss of cortical cytoskeletal stability (Fig. S2A and S2B).

### CD32b is associated with the IgD-BCR nanocluster and stabilizes CD20 expression

The 20-fold upregulation of CD32b on the surface of PKCδ KO Ramos B cells suggests that this inhibitory Fc receptor also has a constitutive gatekeeper function for the resting B cell stage^31^. To test for this, we generated Ramos clones with a CD32b deficiency (CD32b KO) or CD32b overexpression (CD32b-tr). Ramos wt cells express very low amounts of CD32b on their surface, but the CD32b KO line clearly showed that this level can be further reduced to background levels (Fig. 2A). Transfection of Ramos B cells with a human CD32b lentiviral expression vector resulted in a 500-fold increase of CD32b on the surface of CD32b-tr Ramos cells (Fig. 2B). To test for alterations of B cell marker proteins on CD32b-deficient or - overexpressing Ramos cells (CD32-tr), we directly compared these cells with Ramos wt cells in a flow cytometric analysis. Loss of CD32b resulted in a reduced expression of proteins associated with the IgD-class nanocluster such as CD20, CD40, CXCR4, and the IgD-BCR, and a higher expression of CD19 and the IgM-BCR, as well as the B cell activation marker CD69 (Fig. 2C). Notably, CD19 and the IgM-BCR interact with each other on activated B lymphocytes and the expression pattern on CD32b KO Ramos cells suggests that the loss of CD32b results in a partial B cell activation^32^. The comparison of CD32b-tr to Ramos wt cells shows an opposite phenotype, namely the increased expression of proteins associated with the IgD-class nanocluster (Fig. 2D). The effect of the CD32b expression on the stability of the IgD-class nanocluster suggests that CD32b is part of this membrane nanocluster. To test for this, we employed the Fab-based proximity ligation assay (Fab-PLA), which monitors the 10-20 nanometer proximity between the IgD-BCR and CD32b on the cell surface (Fig. 2E and 2F)^33^. The IgD:CD32b PLA signal monitored from Ramos wt is drastically increased upon CD32b overexpression in CD32-tr, reaching a level that is similar to the PLA signal monitored by human healthy donor (HD) B cells. As a negative control for this analysis, we tested Ramos CD32b KO cells, which show no PLA signal. The comparable IgD-BCR/CD32b PLA signal obtained from the analysis of CD32b-tr and HD B cells suggests a similar expression of CD32b on these two cell types. This was verified by a comparative flow cytometry analysis of Ramos CD32b-tr and HD B cells that also confirmed the higher CD20 expression on the Ramos CD32b-tr B cells (Fig. S3). Because of these features and the more stable IgD-class nanocluster we chose CD32b-tr Ramos B cells for our further studies of the cellular function of CD20.

**Figure 2:**
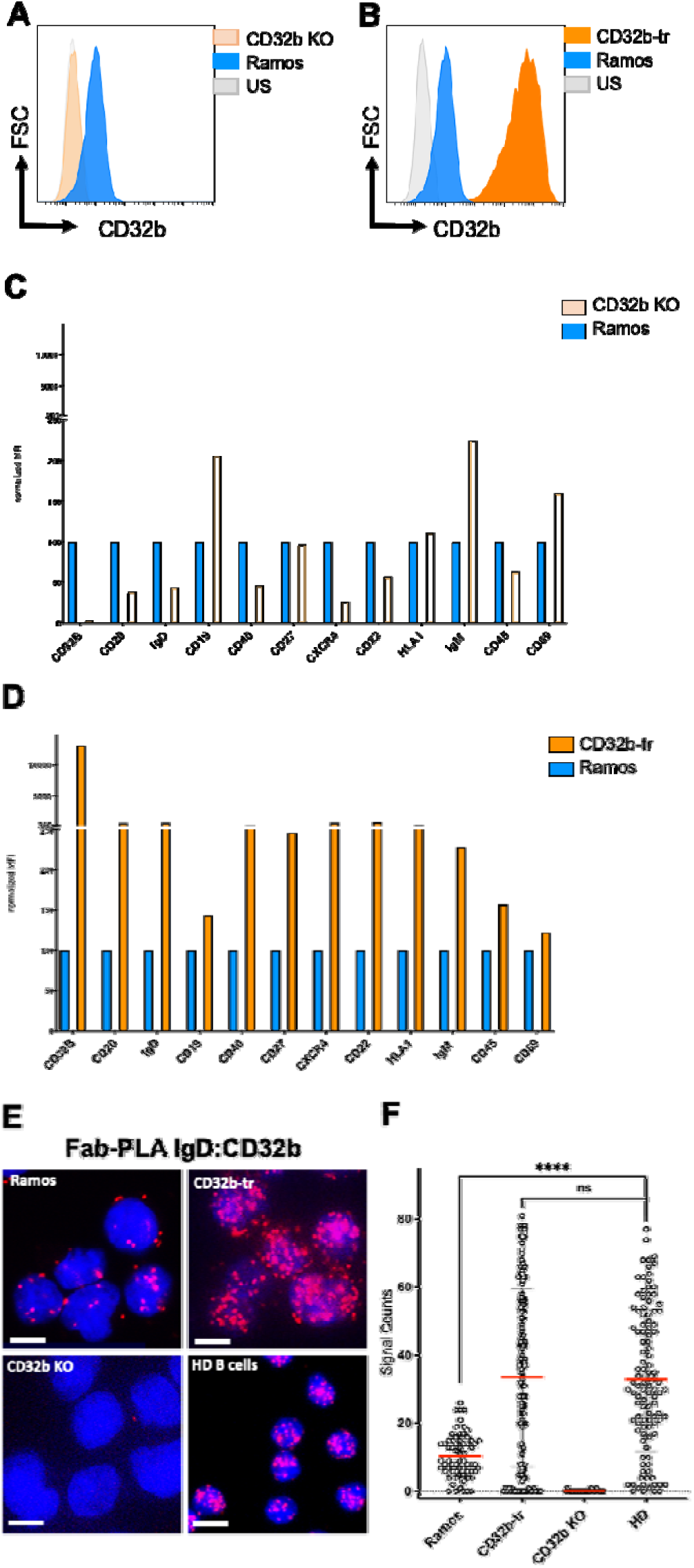
Functional association of the Fc-receptor CD32b with the IgD-BCR nanocluster. **A, B.** Flow cytometry analysis of CD32b expression on the surface of CD32b KO (beige) or CD32b-tr (orange) in comparison to WT Ramos B cells (blue). The unstained control (US) is shown in light grey. **C, D.** Flow cytometry analysis of the expression of B cell marker proteins on the surface of CD32b KO (beige) or CD32b-tr (orange) in comparison to WT Ramos B cells, (blue). Expression data of CD32b KO or CD32b-tr cells are normalized to those of WT Ramos set to 100%. **E.** Fab-PLA study of the IgD-BCR:CD32b proximity on the surface of WT Ramos (upper left), CD32b-tr, (upper right), CD32b KO (lower left), and isolated healthy donor (HD) B cells (lower right). PLA dots in red and nuclei stained blue by DAPI, Scale bar 5 µm, **F.** Quantification of Fab-PLA data with CellProfiler and calculation of significance between the groups with PRISM 10, one-way ANOVA. Students t-test shows no significant difference between CD32-tr and HD B cells. Rea bars show mean values.

### Function of CD20 serine residues and their association with 14-3-3 adaptor proteins

The serine/threonine phosphoproteome analysis of Ramos and its PKCδ-deficient subclones suggests that 4 serine residues in the N-terminal cytosolic tail (S7, S35/36, S49) and 3 serine residues in the C-terminal cytosolic tail (S225, S253/254) of CD20 are constitutively phosphorylated by PKCδ. Interestingly, the three most significant CD20 serine phosphorylation sites (S35/36, S225) are found in the context of an RxxS motif that, once phosphorylated, could become a binding target of the 14-3-3 adaptor family members, which function by binding to serine/threonine-phosphorylated intracellular proteins and altering the conformation, activity, and subcellular localization of their binding partners (Fig. 3A)^34–36^. Additionally, another serine residue (S221) is located near the plasma membrane. Neither the phosphorylated nor the unphosphorylated form of S221 was detected in the phosphoproteome, perhaps because of an adjacent palmitoylation at C220^37^. S221 lies within a classical RxxSxP binding motif for 14-3-3. To eliminate all possibilities of 14-3-3 binding, S221 was also mutated to alanine. To test for the function of the identified serine residues, we generated expression vectors for WT or mutant CD20 proteins carrying a serine to alanine (S/A) exchange of either the 4 N-terminal (Nmut), the 4 C-terminal (Cmut), or all 8 serine residues (NCmut). For purification purposes, the expressed CD20 proteins also carry a C-terminal flag-tag (Fig. 3B). Next, we employed the CRISPR/Cas-9 method to generate CD20-deficient Ramos B cells that were transfected with retroviral expression vectors encoding either CD20 WT or tail mutants (Nmut, Cmut, or NCmut) of CD20. The obtained CD20 transfectants were sorted and further transfected with the human CD32b lentiviral expression vector. Flow cytometry analysis showed that all four double-transfectants expressed CD20 and CD32b at levels similar to the CD32b-tr Ramos line (Fig. 3C). A similar expression was also found for CD19 as well as the IgM- and IgD-BCR. However, in comparison to the CD32b-tr Ramos line and the CD20 WT transfectants, Ramos cells expressing the Nmut, Cmut, or NCmut CD20 mutant displayed a higher expression of the activation marker CD83 suggesting that the S/A exchange partially compromises the gatekeeper function of CD20 (Fig. 3C, far right)^38^.

**Figure 3:**
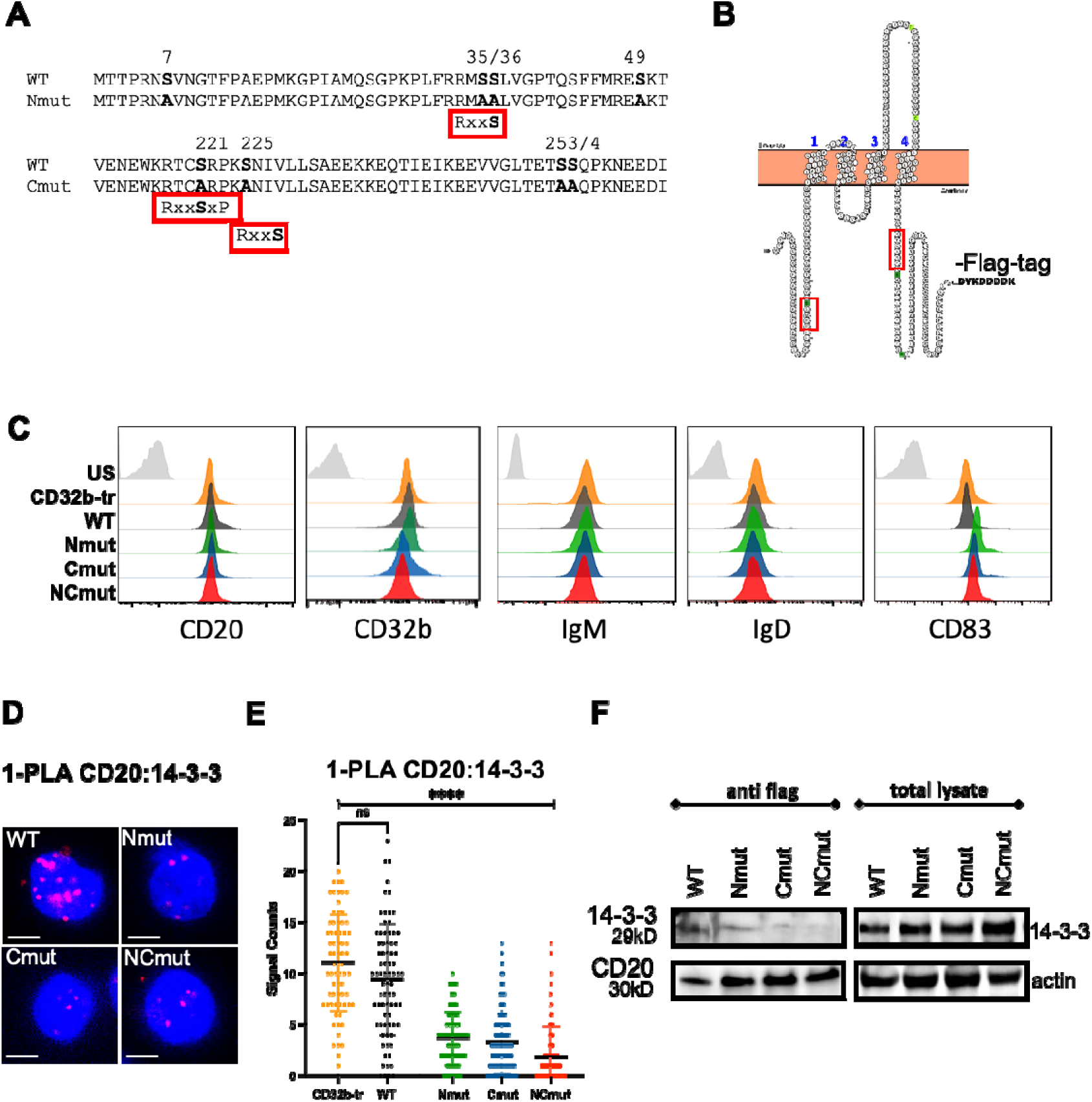
Mutational analysis of CD20 serine residues and their association with 14-3-3. **A.** Protein sequence of the N-terminal (top) and part of the C-terminal cytoplasmic tail of human CD20 (bottom). Below the WT sequence, the sequences with the introduced S/A mutations (bold letters) at the N-terminal (Nmut) and C-terminal (Cmut) cytoplasmic tails of CD20 are shown. The location of the RxxS motifs implicated in 14-3-3 binding is shown below the sequence **B.** Membrane topology of human CD20 with a C-terminal flag-tag (CD20-flag). Red boxes indicate the location of the two RxxS motifs **C**. Flow cytometry analysis of CD32b-tr Ramos cells (orange) and reconstituted Ramos cells expressing CD20 WT (black), Nmut (green), Cmut (dark blue) or NCmut (red). The Ramos cells are tested for the expression of CD20, CD32b, IgM, IgD, and CD83. Unstained (US) CD32-tr cells are shown in grey**. D.** Fab-PLA study of the CD20:14-3-3 proximity in WT (upper left), Nmut (upper right), Cmut (lower left), and NCmut (lower right) Ramos B cells. PLA dots in red and nuclei stained blue by DAPI, Scale bar 5 µm. **E**. Quantification of PLA data by CellProfiler. Significant differences between groups were calculated with PRISM10, one-way ANOVA. Students t-test shows no significant difference between CD32-tr and WT Ramos cells. **F.** Western blot analysis of the CD20-flag/14-3-3 association in WT, Nmut, Cmut and NCmut Ramos B cells. The amount of 14-3-3 protein in anti-flag immunoprecipitates and total lysates is show on the left and right, respectively. Lysates were taken from at least 3 independently generated Ramos cell lines.

The finding that 4 of the 8 serine residues under study are part of an RxxS motif sequence suggests that CD20 is not only constitutively phosphorylated by PKCδ but also constitutively associated with a 14-3-3 adaptor protein. To test for this, we used the 1-PLA assay to monitor the colocalization of CD20 with a 14-3-3 adaptor protein in CD32b-tr Ramos cells and CD32b/CD20 double-transfectants expressing in addition to CD32b, either CD20 WT or one of three different tail mutants of CD20 (Fig. 3D and 3E). The 1-PLA assay employs CD20 and 14-3-3 specific DNA-coupled antibodies that detect a colocalization of the two proteins in the 10-40 nanometer range^33^. A colocalization of CD20 with 14-3-3 is detected in the CD20 WT, but to a lesser extent in the CD20 Nmut, Cmut or NCmut Ramos cells. Western blot analysis of anti-flag immunoprecipitates of Ramos cell lysates confirms that, compared to CD20 WT, the S/A mutants of CD20 are less well associated with the 29 kD 14-3-3 protein (Fig. 3F).

### The GEF-H1 activity is regulated by the cytoplasmic tails of CD20

14-3-3 is a dimeric adaptor protein that can stabilize a CD20 homodimer and/or the heterodimeric association of CD20 with other RxxS motif-containing proteins. In a preliminary study, we found that exposure of B cells to anti-CD20 antibodies activates the RhoA/Rock1 signaling pathway (see below). A key activator of this pathway is the GDP/GTP exchange protein GEF-H1 (also known as ARHGEF2), which is constitutively associated with the MT network in resting B cells^39–44^. The intrinsically disordered regions in the C-terminal tail of GEF-H1^40^ carries an RRxxR and an RxxSxP motif (Fig. 4A). The RRxxR motif is a known binding target for light chains of the dynein motor complex (such as LC8 or Tctex-1) and is thought to be one of the interfaces linking GEF-H1 to MTs^42^. The serine residue S886 within the RxxSxP motif of GEF-H1 is phosphorylated by the p21-activated kinase-2 (PAK2), a modification that is known to inhibit GEF-H1 activity^45^. Binding of a 14-3-3 adaptor protein to the phosphorylated RxxSxP motif may connect GEF-H1 to the constitutively phosphorylated CD20 tails. To test for this, we used the 1-PLA assay and found a colocalization of CD20 and GEF-H1 in the CD20 WT transfectant and to a lesser extent in the CD20 Nmut, Cmut or NCmut transfected Ramos cells (Fig. 4B and 4C). Western blot analysis of anti-flag immunoprecipitates of CD20 proteins in the Ramos cell lysates confirmed that the tail mutants of CD20 are less well associated with GEF-H1 than CD20 WT (Fig. 4D). In particular, the NC double-mutant of CD20 is hardly associated with GEF-H1, compared to CD20 WT.

**Figure 4:**
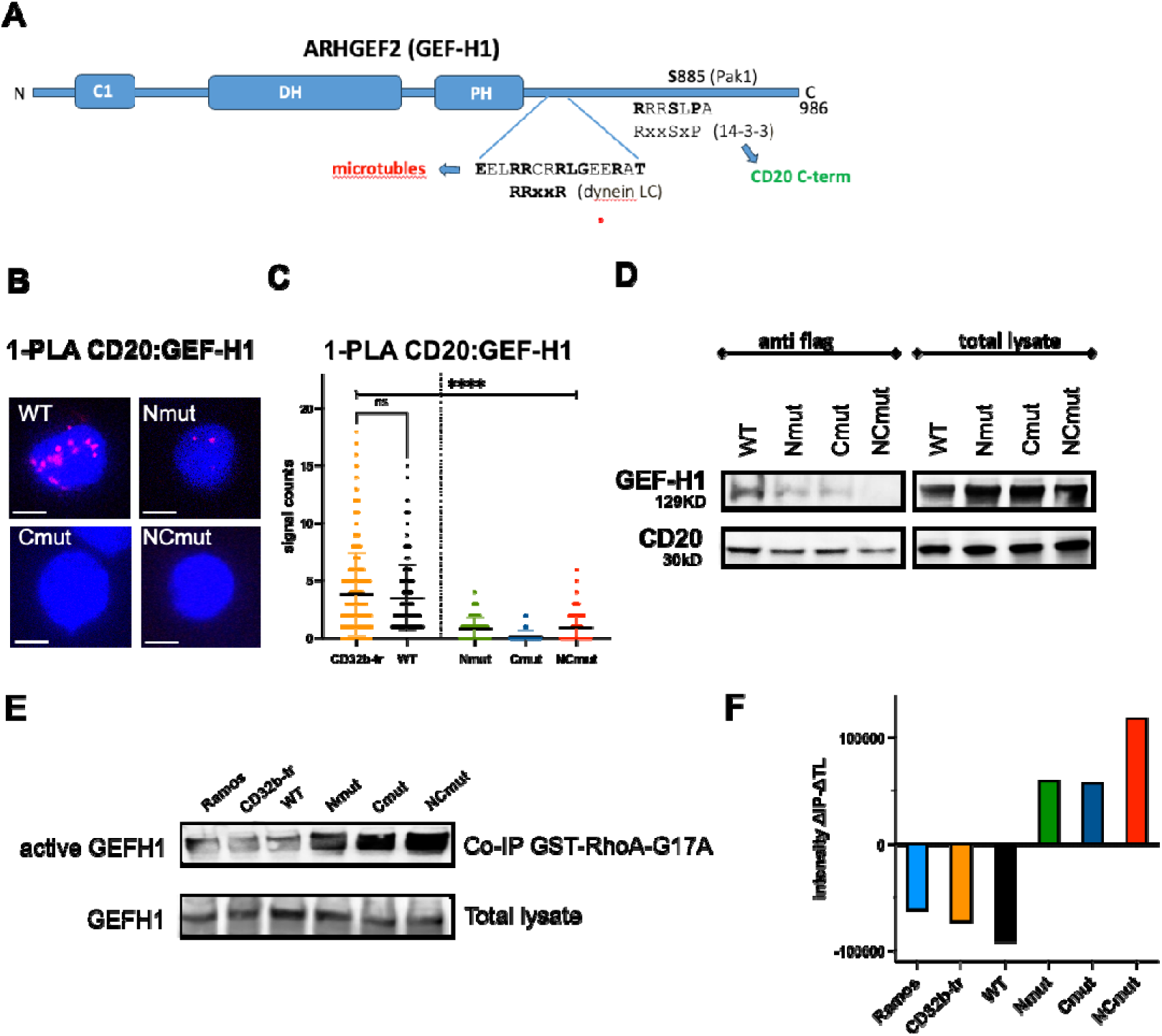
Constitutive association of CD20 with the GDP/GTP exchange factor GEF-H1. **A.** Schematic drawing of the domain structure of the RhoA guanine nucleotide exchange factor GEF-H1 (ARHGEF2). The functional domains are drawn as blue boxes. The location of the RRxxR motif implicated in dynein light chain binding, and the Pak2 phosphorylated S886 within the 14-3-3 binding **R**xx**S**x**P** motif is indicated. **B**. 1-PLA study of the CD20: GEF-H1 proximity in WT (upper left), Nmut (upper right), Cmut (lower left), and NCmut (lower right) Ramos B cells. PLA dots in red and nuclei stained blue by DAPI, Scale bar 5 µm. **C**. Quantification of PLA of B. with CellProfiler. Significant differences between groups were calculated with PRISM10, one-way ANOVA. Students t-test shows no significant difference between CD32-tr and WT Ramos cells. **D.** Western blot analysis of the constitutive CD20-flag/GEF-H1 association in WT, Nmut, Cmut and NCmut Ramos B cells. The amount of GEF-H1 protein in the anti-flag immunoprecipitates and total lysates is show on the left and right, respectively. **E**. Western blot analysis of free GEF-H1 in the lysates of Ramos and CD32b-tr cells as well as reconstituted Ramos cells expressing CD20 WT, Nmut, Cmut or NCmut. Amount of GEF-H1 purified with GST-RhoAG17A beads and in total lysate is shown at the top and bottom, respectively. **F.** Quantification of the Western blot data with Image Studio Lite (Licor) showing the difference of signal intensity in the immunoprecipitates (IP) versus total lysates (TL).

The amount of the GDP/GTP exchange active GEF-H1 protein in the cytosol of Ramos B cells can be tested by a pulldown assay employing a G17A trapping mutant of RhoA, which stabilizes the intermediate RhoA/GEF-H1 interaction^46^. For this, lysates of the different Ramos cell lines were incubated with GST-RhoA-G17A-coupled GST beads, and the bound GEF-H1 was monitored by western blotting. Compared to CD20 WT, the cytosol of Ramos cells expressing the Nmut, Cmut or NCmut contains a higher amount of active GEF-H1/RhoA complexes (Fig. 4E and 4F). Taken together, these data suggest that the constitutively phosphorylated serine residues within the cytosolic tails of CD20 regulate the RhoA/Rock-1 pathway via the binding and sequestering of GEF-H1.

### Exposure to anti-CD20 antibody activates the RhoA/Rock-1 pathway

We next asked whether or not the exposure of Ramos B cells to a therapeutic anti-CD20 antibody such as rituximab (RTX) alters the interaction of CD20 with components of the RhoA/Rock-1 pathway. For this we employed the 1-PLA assays to monitor changes in the proximity of CD20 to 14-3-3, GEF-H1, RhoA and Rock-1 in Ramos CD32b-tr as well as CD20 WT or CD20 NCmut transfectants before or after a 5 min exposure to RTX (Fig. 5). The RTX treatment reduced the CD20/14-3-3 proximity in CD32b-tr and CD20 WT transfected Ramos cells to the level of unstimulated NCmut Ramos cells (Fig. 5A and 5B) and the same was found for the CD20/GEF-H1 proximity (Fig. 5C and 5D). Interestingly, the 1-PLA assays show an increased proximity of CD20 and RhoA (Fig. 5E and 5F) as well as CD20 and Rock-1 in RTX-treated Ramos cells expressing CD20 WT but not those expressing CD20 NCmut. This study suggests that in RTX-treated B cells, CD20 stabilizes a RhoA/Rock-1 complex at the inner leaflet of the plasma membrane in a 14-3-3-dependent manner^47,48^. Importantly, the C-terminus of Rock-1 also carries an evolutionary conserved RxxS motif. Thus, it is feasible that CD20 functions not only as a gatekeeper of the resting state, but also as a switch that promotes activation of the RhoA/Rock-1 pathway in antigen- or RTX-exposed B cells. As a control, we determined whether active GEF-H1 is formed after RTX activation following GDP/GTP exchange. The pull-down assay with the G17A trapping mutant of RhoA shows an increase in active GEF-H1/RhoA complexes, especially in Ramos, CD32b-tr and WT cells (Fig. S4A and 4B).

**Figure 5:**
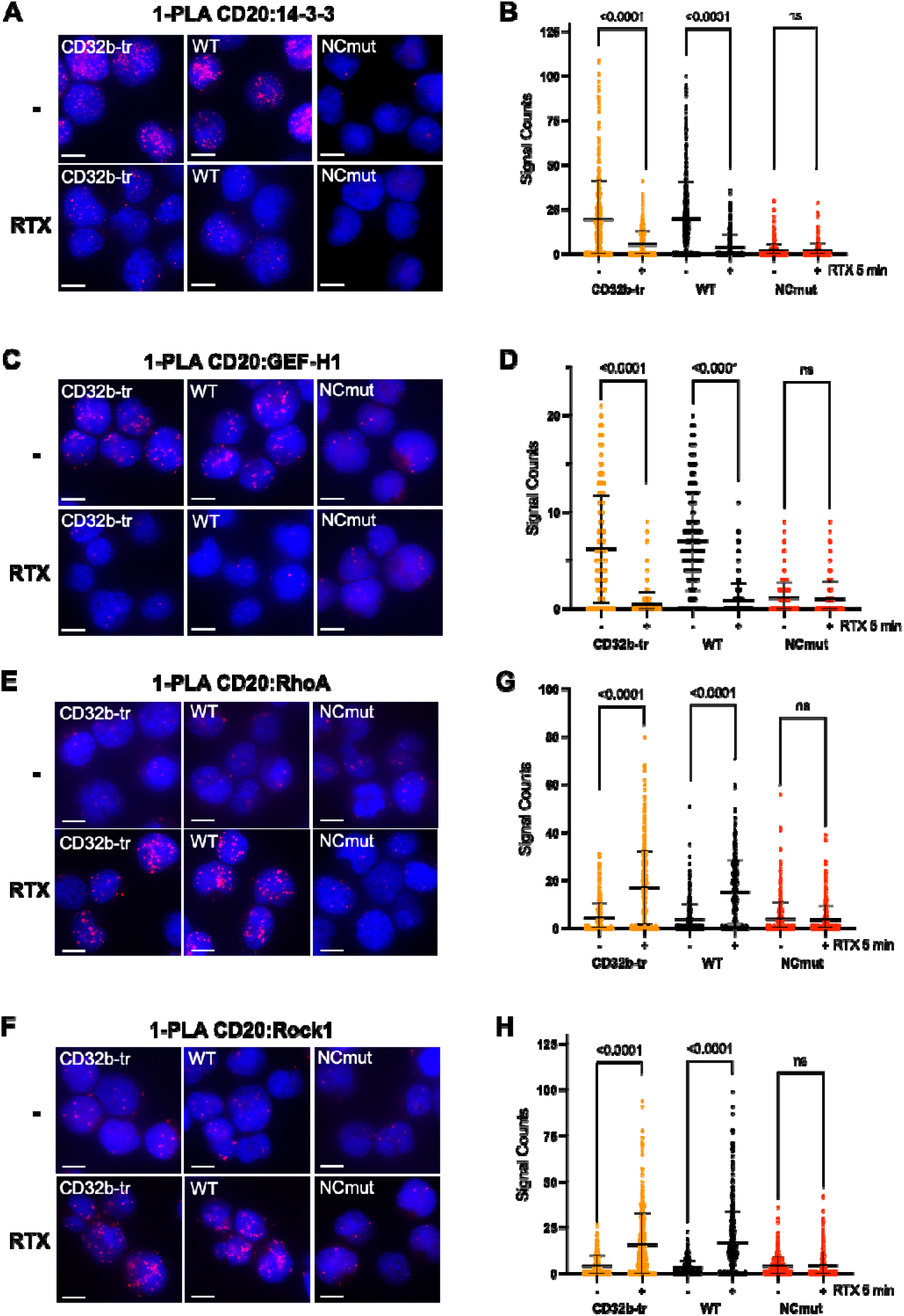
Loss of GEF-H1 and gain of the RhoA/Rock1 association of CD20 upon RTX treatment. 1-PLA study of **A.** the CD20:14-3-3, **C.** the CD20:GEF-H1, **E.** the CD20:RhoA and **F.** the CD20:ROCK1 proximity in CD32b-tr Ramos cells (left) and reconstituted Ramos cells expressing WT (middle), or NCmut CD20 (right). The studied Ramos cells were either untreated (top) or exposed to RTX for 5 min (bottom). PLA dots in red and nuclei stained in blue by DAPI, Scale bar 5 µm. **B**, **D, G, H.** Quantification of the 1-PLA data with CellProfiler. Significant differences between two groups were calculated with PRISM10 Students t-test.

We then used the phosphoflow assays to monitor alteration in the phosphorylation of members of the RhoA/Rock-1 pathway after the RTX treatment of CD32b-tr, CD20 WT or NCmut Ramos cells (Fig. S5). Exposure of CD32b-tr or CD20 WT Ramos cells to RTX results in a reduction of the S886 phosphorylation of GEF-H1 and the loss of the inhibitory S188 phosphorylation of RhoA (Fig. S5A and 5B). The same treatment resulted in an increased phosphorylation of tyrosine Y914 of Rock-1, a modification that is associated with Rock-1 activation (Fig. S5C)^49,50^. Importantly, all these amino acids are less well phosphorylated in the NCmut Ramos cells and their phosphorylation state is not significantly altered after exposure of the cells to RTX. Together these data demonstrate that the cytosolic serine/threonine residues studied connect CD20 to the RhoA/Rock-1 signaling pathway.

Exposure to the therapeutic antibody RTX activates the RhoA/Rock-1 signaling pathway not only in Ramos but also naïve human B cells. This is indicated by the time-dependent alteration of the phosphorylation status of GEF-H1, RhoA and Rock-1 upon the treatment of HD B cells with 10 ug/mL of RTX (Fig. S5D-5F).

The 1-PLA assays of HD B cells before or after a 5 min exposure to RTX show a reduction of the 14-3-3 (Fig. 6A and 6B) and GEF-H1 (Fig. 6C and 6D) proximity to CD20 and at the same time a strong recruitment of RhoA (Fig. 6E and 6F) and Rock-1 (Fig. 6G and 6H) in the vicinity of CD20 in the RTX-treated cells. Thus, the CD20 engagement not only liberates and activates GEF-H1 but also results in an increased coupling of the engaged CD20 to the RhoA/Rock-1 signaling module. Interestingly not only the RTX treatment but also exposure of HD B cells to a MT polymerization inhibitor such as nocodazole (Noc) changes the CD20 association with components of the RhoA/Rock-1 signaling pathway (Fig. 6E-6H).

**Figure 6:**
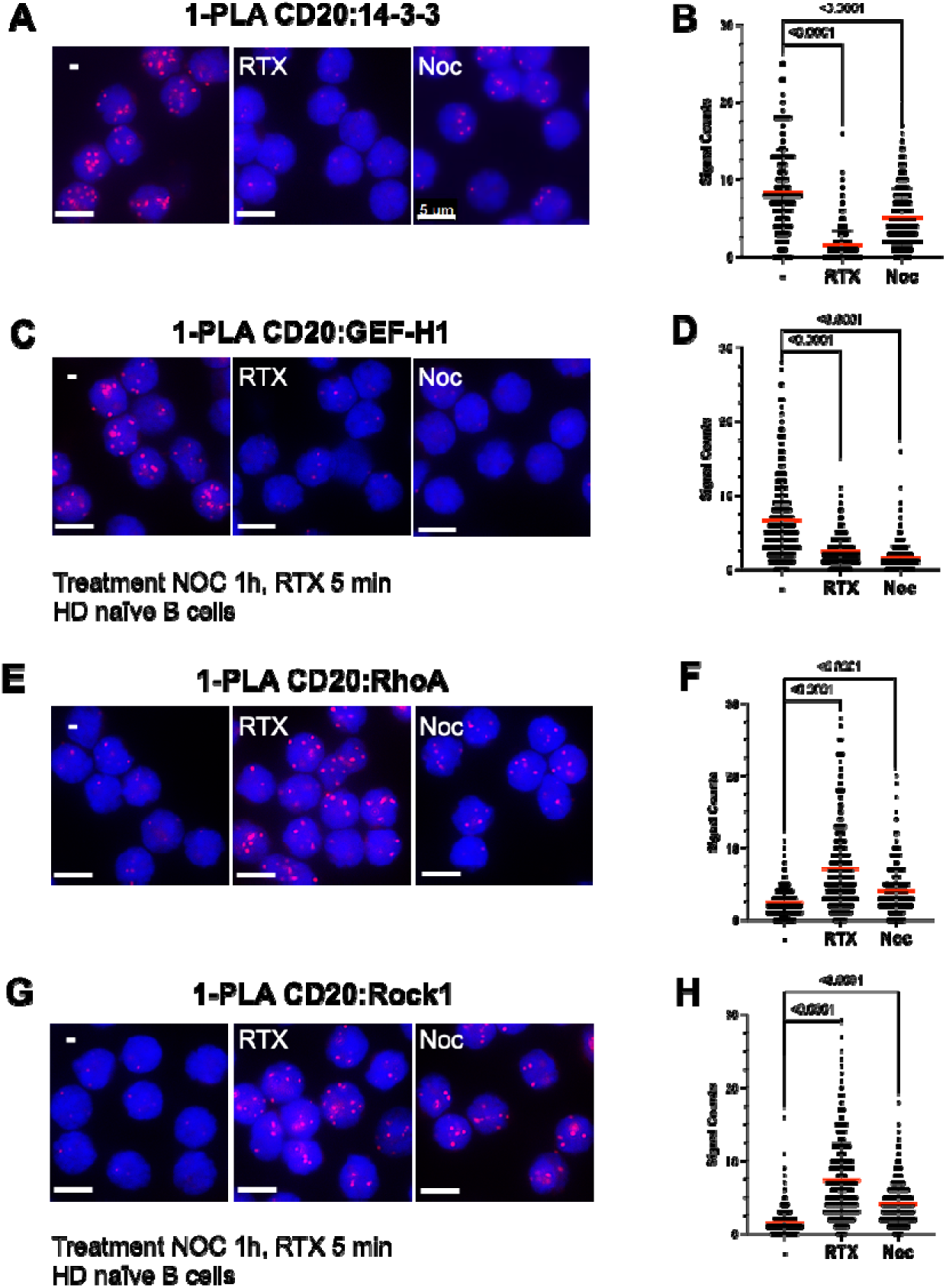
Altered CD20 association in RTX or nocodazole treated HD B cells. 1-PLA study of **A.** the CD20:14-3-3, **C.** the CD20:GEF-H1, **E.** the CD20:RhoA and **F.** the CD20:ROCK1 proximity in HD B cells unstimulated (left), RTX-exposed for 5 min (middle), or treated for 1 h with Noc (right). PLA dots in red and nuclei stained in blue by DAPI, Scale bar 5 µm. **B**, **D, G, H.** Quantification of 1-PLA data with CellProfiler. Significant differences between two groups were calculated with PRISM 10 Students t-test.

### Destabilization of MTs and the IgD-BCR nanocluster by anti-CD20 antibodies

It is well known that GEF-H1 is associated with the MT network and that MT destabilization results of GEF-H1 liberation and increased activity^39,41–43,51,52^. The result of our 1-PLA assay (see above) suggests that the activation of RhoA/Rock-1 signaling pathway by RTX is mediated by the destabilization of the MT cytoskeleton. To test for this, we pretreated HD B cells for 1 hour with either Noc or with the MT stabilizer docetaxel (Tax) and then measured RTX-induced changes by flow cytometry (Fig. 7A and 7B). The RTX-induced reduction of CD20 abundance and the increased Rock1 phosphorylation can be prevented by pretreating the HD B cells for 1 hour with Tax, whereas the Noc-pretreated HD B cells show already an Rock1 activation with no further increase by RTX. Thus, the immediate effect of the binding of RTX to CD20 seems to be the destabilization of the MT network resulting in GEF-H1 liberation and the activation of the RhoA/Rock-1 signaling pathway.

**Figure 7:**
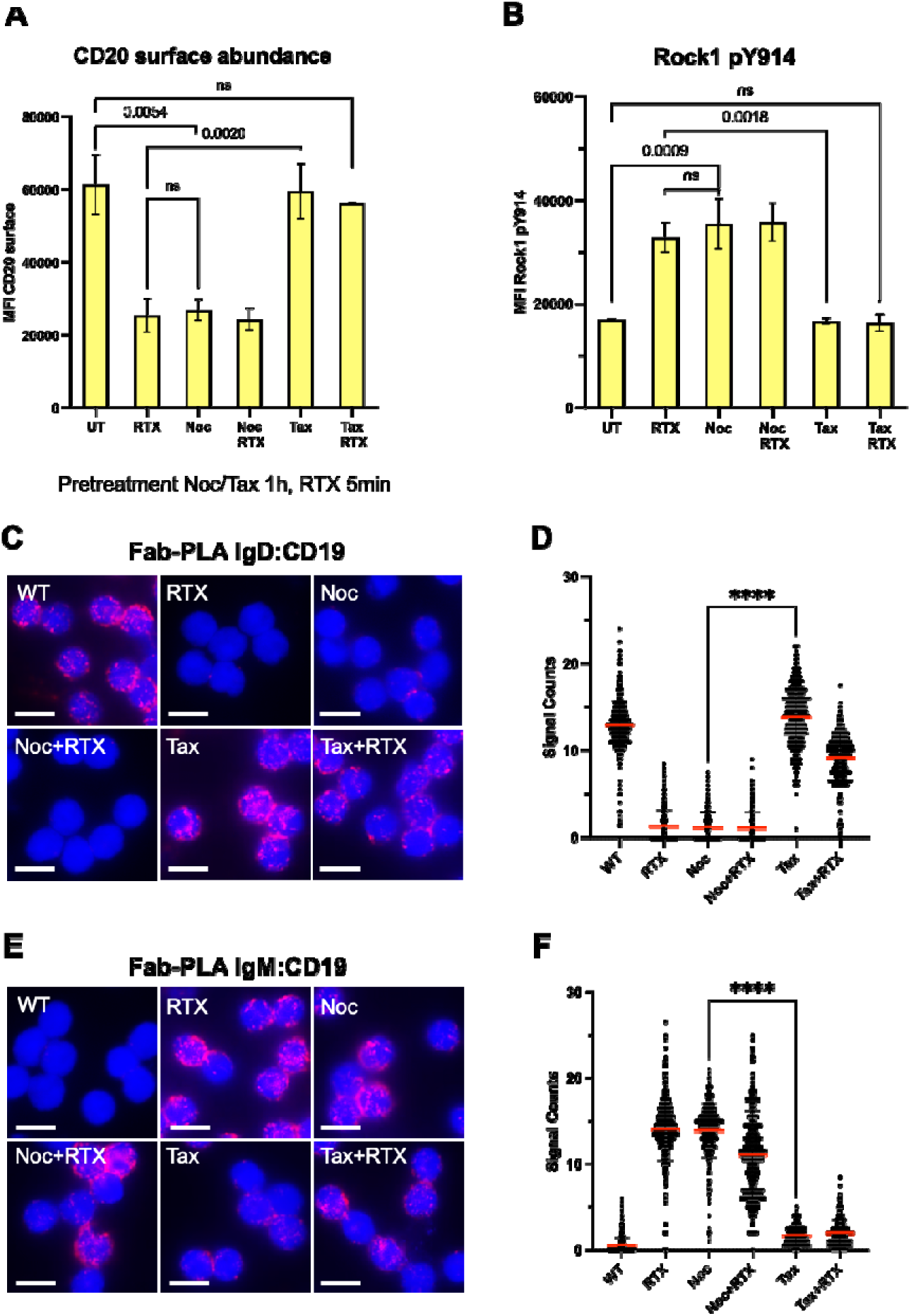
Stabilization of the IgD-BCR nanocluster and CD20 by the MT network. **A**. CD20 expression on the surface of untreated (UT) HD B cells or after RTX-stimulation for 5 min. As indicated, some of these HD B cells were pretreated for 1 h with either nocodazole (Noc) or docetaxel (Tax). Shown is a summary of the MFI data of >3 flow cytometry experiments. PRISM10 students t-tests were used to calculated the significant differences between the experimental groups. **B**. Y914 phosphorylation and activation of Rock1 in untreated (UT) HD B cells or B cells stimulated for 5 min with RTX. As indicated, HD B cells were kept untreated, or pretreated for 1 h with nocodazole (Noc), or docetaxel (Tax). A summary of the MFI data of >3 flow cytometry experiments is presented. Students t-tests were used to calculated the significant differences between the groups. Fab-PLA study of **C.** the IgD-BCR:CD19 or **E.** the IgM-BCR:CD19 proximity of HD B cells, kept untreated or stimulated for 5 min with RTX without or with pretreatment of the B cells with either Noc or Tax for 1h as indicated. PLA dots in red and nuclei stained blue by DAPI, Scale bar 5 µm. **D, F.** Quantification of Fab-PLA data with CellProfiler. Significant differences between groups were calculated with PRISM 10 Students t-test.

MT dynamics are tightly regulated. To test whether binding of RTX to CD20 can influence or induce the destabilization of MTs, Ramos CD32b-tr cells were treated with RTX and labeled with mouse anti-alpha-tubulin to visualize tubulin structures by immunofluorescence microscopy. In the Airyscan Super-Resolution (SR) imaging results (Fig. S6) we could visualize the peeling of MT polymers, the outwardly curved spiral protofilaments at the end of shortening MT upon RTX exposure (Fig. S6A and B). In a second approach with HD naïve B cells (Fig. S7A-C), Airyscan (SR) confocal images of the labeled α-tubulin showed a spatial distribution of small “ram’s-horn” MT intermediates upon RTX treatment (Fig. S7B)^53,54^. Interestingly, a pretreatment with Tax before RTX exposure could stabilize the B cell integrity and reduce MT depolymerization (Fig. S7C).

On resting B cells CD20 and CD19 reside in the same protein/lipid nanocluster in close proximity to the IgD-BCR, and the exposure to RTX leads to IgD-BCR/CD19 dissociation and the formation of an IgM-BCR/CD19 complex^26^. Taking advantage of the Fab-PLA technique, we next asked whether or not the MT network is involved in the nanoscale receptor organization and reformation. Indeed, the exposure of the HD B cells to either RTX or Noc results in a dissolution of the IgD-BCR nanocluster and IgM-BCR/CD19 conjugation. The RTX-induced nanoscale receptor rearrangements are partially prevented by the pretreatment of the HD B cells with Tax (Fig. 7C-7F). Taken together, these data show that in resting B cells the nanocluster of the IgD-BCR is connected to the MT network and that the cytosolic tails of CD20 play an important role in this connection and its formation.

## Discussion

### PKCδ and CD20: a kinase/substrate pair with a gatekeeper function

We previously have shown that CD20 is part of the membrane nanocluster of the IgD-BCR and acts as a gatekeeper for the resting state of B lymphocytes^26^. Here we describe that serine residues within the cytoplasmic tails of CD20 play an important role for this gatekeeper function. Specifically, we show that 6 serine residues of CD20 are constitutively phosphorylated by the serine/threonine kinase PKCδ and that this modification couples CD20 to the MT network in resting and the RhoA/ROCK1 signaling pathway in activated B cells. The exchange of the serine to alanine residues did not alter the amount of CD20 on the B cell surface, but it has an impact on the B cell activation state as the CD20 mutated Ramos cells displayed a higher expression of the CD83 activation marker^38^. These data suggest that the phosphorylated serine residues are connected to the gatekeeper function of CD20. The notion that PKCδ also acts as a gatekeeper for the resting state of B lymphocytes is supported by several genetic data^27,29,55,56^. The PKCδ KO mice develop autoimmunity^57^ and loss-of-function mutations of PKCδ in humans are associated with autoimmune diseases such as systemic lupus erythematosus (SLE) or a B cell lymphoproliferative syndrome^58^. Our finding that CD20 is a substrate of PKCδ provides a novel mechanistic insight in the development of these autoimmune diseases.

PKCδ has multiple substrates at the membrane, in the cytosol and inside the nucleus^55^. Thus, the targeting of PKCδ to a specific subcellular location may determine its biological function in different settings. Like many other kinases, PKCδ is regulated by an autoinhibitory mechanism where the N-terminal regulatory domain is folded over the C-terminal catalytic domain, thus inhibiting the kinase activity. The specific targeting of PKCδ is connected with an opening of the autoinhibited conformation and kinase activation. Interestingly, the SRC-family kinase Lyn can phosphorylate and bind to tyrosine 52 of PKCδ^59,60^. The formation of the Lyn/PKCδ complex activates both kinases and places them at the inner leaflet of the plasma membrane. We have previously found that, after an Igα phosphorylation, the IgD-BCR is associated with PKCδ^24^. It is thus feasible that an active Lyn/PKCδ complex is formed within the IgD-BCR nanocluster, thereby explaining the constitutive serine phosphorylation of CD20 at this location.

### CD20 binds GEF-H1 and inhibits the GDP/GTP exchange activity in resting B cells

Four of the 8 mutated serine residues of CD20 are part of conserved RxxS or RxxSxP motifs that are well-known binding sites for 14-3-3 adaptor proteins^61^. A constitutive CD20/14-3-3 association in resting human B cells was confirmed by a 1-PLA and immune precipitation. The RxxS and RxxSxP sequence is localized at the CD20 N-terminus and C-terminus, respectively. The binding of the dimeric 14-3-3 adaptor to these sequences can either alter the intramolecular conformation of the two cytosolic tails of CD20 or stabilize the formation of a CD20 homodimer. Alternatively, the 14-3-3 dimer could also connect CD20 to other signaling molecules^62,63^. Using a 1-PLA and immune precipitation assays we confirmed that the GDP/GTP exchange factor GEF-H1, which carries a conserved RxxSxP motif at its unstructured C-terminus, is one of the binding partners of the constitutively phosphorylated CD20. The serine residue S886 within the conserved RxxSxP motif of GEF-H1 is constitutively phosphorylated by Pak2, a modification that is associated with the inhibition of the GDP/GTP exchange activity^64^. Thus, two serine/threonine kinases, PKCδ and Pak2, control the formation and silencing of the CD20/GEF-H1 complex.

The activity of GEF-H1 is regulated by the cytoskeleton^52^. Specifically, GEF-H1 is intimately associated with the MT network and this association inhibits its GDP/GTP exchange activity. There are two (non-mutually exclusive) mechanisms how MT binding inhibits GEF-H1. In the first one, polymerized MT stabilize the autoinhibited conformation in which the N-terminal C1 domain is folded over the PH domain of GEF-H1^65^. The second mechanism involves the binding of the light chain protein Tctex-1 of the dynein motor complex to the RRxxR binding motif at the unstructured C-terminal tail of GEF-H1^66,67^. In this way the dimeric dynein motor complex couples GEF-H1 to polymerized MT. The RxxSxP motif that connects GEF-H1 to the phosphorylated CD20 tails lies 60 amino acids downstream of the Tctex-1 binding site. Thus, in resting B cells the autoinhibited GDP/GTP exchange factor functions as an adaptor that connects CD20 to the MT network. Based on the finding that anti-CD20 antibodies can modulate the calcium response in B lymphocytes, it has previously been suggested that CD20 acts as a calcium channel^68,69^. This suggestion is not supported by the CD20 structure which did not reveal a calcium permeable pore^10^. However, we noticed that the N-terminus of the store-operated calcium channel ORAI 1 carries an RRxxR and an RxxS motif^70^. Thus, Tctex-1 and the 14-3-3 adaptor could couple CD20 to the ORAI1 channel and this connection could explain the observed effect of anti-CD20 antibodies on the calcium response in B lymphocytes.

### CD20 can orchestrate the MT-to-actin switch during B cell activation

In migrating, polarized B lymphocytes, the MT organization center (MTOC) is found in the vicinity of the trailing edge and the MT network is involved in the organization of the uropod at this location^71^. We have found that in polarized B cells the IgD-BCR nanocluster is localized at the uropod whereas at the leading-edge the IgM-BCR is associated with microvilli structures that are formed by core bundles of actin filaments^72^. The interaction of CD20 to the MT network may help to stabilize the IgD-BCR nanocluster^73^. Interestingly, apart from the CD20/GEF-H1 complex the IgD-BCR nanocluster harbors several receptors that carry in their cytosolic juxtamembrane region a RRxxR (BAFF-R) or a related RRxxK motif (CD19) for Tctex-1 binding. Thus, multiple contact points to the MT network may promote the stability of the IgD-BCR nanocluster^74^.

In polarized lymphocytes RhoA and ROCK1 are also found at the uropod and a RhoA deficiency or an inhibition of ROCK1 kinase activity prevents the formation of this structure^71,75^. RhoA undergoes geranylgeranylation at its C-terminal CAAX motif and thus is constitutively bound via its lipid anchor to the inner leaflet of the plasma membrane^76^. In spite of its permanent membrane association, RhoA in resting B cells is not found in the vicinity of CD20 and the IgD-BCR nanocluster as the PLA has shown. Only upon MT destabilization and the following dissociation of the GEF-H1/CD20 complex RhoA and ROCK1 are found in the vicinity of CD20. Interestingly, ROCK1 carries a RxxS motif close to its C-terminus and it is feasible that the GEF-H1/CD20 complex in resting B cells is replaced by a RhoA-GTP/ROCK1/CD20 complex in activated B cells. The GEF-H1 activation and liberation from the MT network could have a dual impact on ROCK1 activity and membrane location. The increased GDP/GTP exchange activity of GEF-H1 generates RhoA-GTP that attaches to the RhoA binding domain of ROCK1 and the GEF-H1-deprived CD20/14-3-3 complex is free to bind to ROCK1. Both binding events counteract ROCK1 autoinhibition and localizes the active ROCK1 at the IgD-BCR nanocluster where ROCK1 phosphorylates the myosin light chain (among other substrates), resulting in the formation of myosin-II filaments and actomyosin stress fibers with increased contractility^77^. The stress fiber formation in polarized, antigen-exposed activated B cells^62,78^ may be associated with uropod retraction, the dissociation of the IgD-BCR nanocluster and the observed movement of CD20 and CD19 to the IgM-BCR. According to this scenario CD20 is not only a gatekeeper of the resting state but also an actomyosin bundles associated transporter that promotes the immunoreceptor coupling and organization motif (ICOM)-dependent conjugation of the IgM-BCR with the CD19 coreceptors in activated B lymphocytes^79^.

### The effect of anti-CD20 antibodies on cytoskeleton reorganization

Previously it was thought that binding of anti-CD20 antibodies did not have a direct effect on the physiology of targeted B cells, but simply provided a flag for their destruction via phagocytosis or ADCC mechanisms^80^. However, several studies indicated that anti-CD20 antibodies induce an apoptosis program in the targeted B cells^81^. We found that exposing B cells to anti-CD20 antibodies results in the dissolution of MTs and MT instability, which is associated with apoptosis ^43,49,82^. Furthermore, caspase 3, an important effector of the apoptosis program, can cleave and activate ROCK1^49^. Interestingly, it is the cleaved, active ROCK1 fragment, which carries the C-terminal RxxS motif and therefore could still be associated with CD20. In this way the active kinase domain of ROCK1 remains bound to the plasma membrane in a RhoA-GTP independent manner. A recent 3-dimensional super-resolution study has localized the RTX-engaged CD20 on actin bundles within microvilli structures^83,84^. This is in line with our findings that the IgM-BCR is associated with an actin-based network of ridges of microvilli structures and that CD20 is found in the vicinity of the IgM BCR after exposure to either cognate antigen or anti-CD20 antibodies^72^. Together these data support the assignment of CD20 as a switch factor, flexible to interact with the MT and alternatively the actomyosin cytoskeleton in resting and activated B lymphocytes, respectively.

The finding that the exposure to anti-CD20 antibodies results in dissolution of MT bundles and protofilaments and that MT-destabilizing drugs can mimic the effect of an anti-CD20 treatment of B lymphocytes not only gives a new insight in the therapeutic effect of the anti-CD20 therapy but also could have an impact on the improvement of such a therapy. It is known that anti-CD20 treatment results in the reduction of the amount of CD20 on the B cell surface and this could reduce the targeting efficiency^85,86^. The exposure of healthy donor B cells to MT stabilizing drugs such a Taxol derivates could prevent the downregulation of antibody-bound CD20 and thus increase the B cell targeting function of the anti-CD20 therapy. However, there also exist data showing that the ADCC-dependent killing of anti-CD20 labelled B cells by NK cells requires dynamic alterations of the MT network. If this is the case, a combination of anti-CD20 therapy with MT-destabilizing drugs such a vincristine could enhance the NK-mediated cytotoxicity of anti-CD20-targeted B cells^87^. While these potential modification of the anti-CD20 therapy requires more studies, the cellular function of CD20 as an adaptor between antigen receptors and the cytoskeleton is now established and CD20 is no longer an enigma of B cell biology.

## Author-contribution

K.K. conceptualization; data curation; formal analysis; funding acquisition; investigation; methodology; project administration; resources; supervision; validation; visualization; writing – original draft; writing – review and editing. C.E.L. conceptualization; data curation; formal analysis; investigation; methodology; project administration; resources; validation; J.B., G.A., A.N., and C.M conceptualization, data curation; formal analysis; investigation; methodology; project administration; resources; software; validation; visualization; writing – review and editing. L.R. conceptualization; data curation; formal analysis; investigation; methodology; resources; software; validation; visualization; writing – review and editing. N.H. conceptualization; data curation; investigation; methodology. R.N. data curation; formal analysis; funding acquisition; investigation; methodology; resources; software; supervision; validation; visualization; writing – review and editing. R.E.V., B.W., and K.W. funding acquisition; investigation; resources; validation; writing – review and editing. M.R. conceptualization; funding acquisition; investigation; methodology; project administration; supervision; writing – original draft; writing – review and editing.

## Supporting information

reagents and Tools

## Acknowledgments

We thank Dr. Christian Klein and the Roche Innovation Center Zurich for financial support and anti-CD20 antibodies and Falk Nimmerjahn, University of Erlangen for the CD32B plasmid. We would like to thank Dr. Baerbel Keller for her help with the RNA sequencing. We thank Dr. Lise Leclercq for correcting the manuscript.

## Supplements

**Supplementary figure 1:**
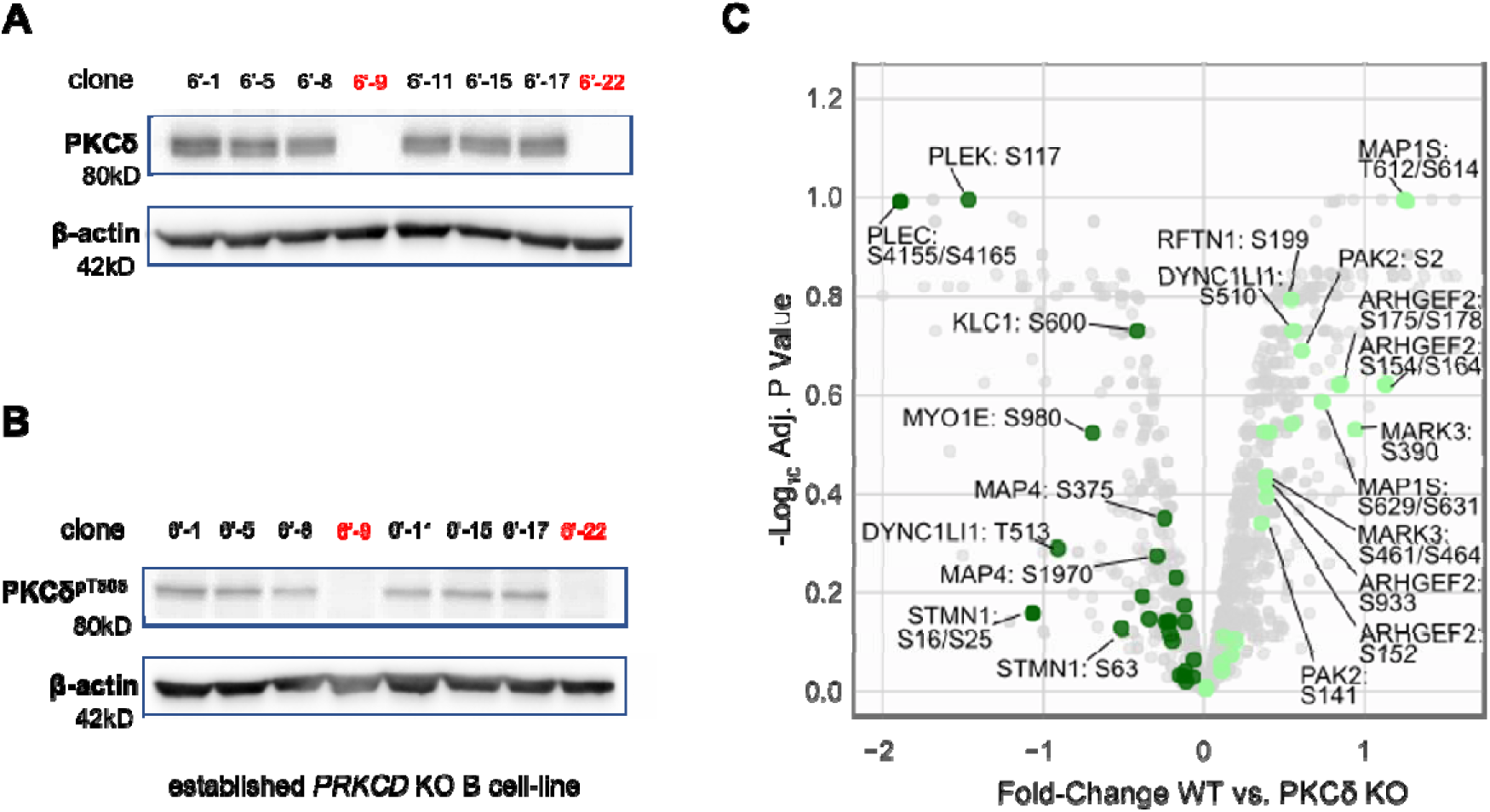
Western blot analysis of PKCδ after CRISPR-Cas9 mediated PKCδ KO. Lysates were stained for PKCδ **A**. or PKCδ phospho-threonine 505 (PKCδ pT505) **B**. Clones 6’-9 and 6’-22 were selected for further PKCδ KO studies. **C**. Volcano blot indicating the changes of the constitutive serine or threonine protein phosphosites of PKCδ KO compared to Ramos cells. On the X-axis the plot depicts log2 fold change of phosphorylated serine or threonine residues in the whole proteome of the two cells lines. The Y-axis shows the log10 FDR adjusted p-value. Decreased phosphorylation of serine residues of the MT pathway are shown in dark green, increase in light green. The phosphorylated serine residues of MT proteins are indicated.

**Supplementary figure 2:**
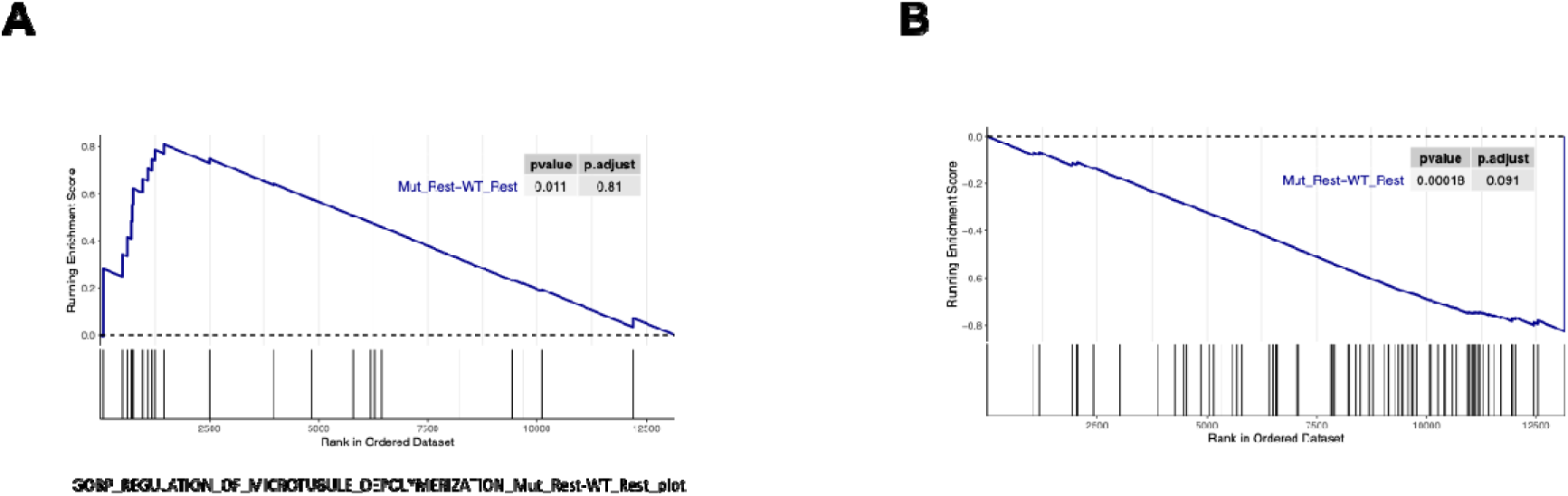
Gene set enrichment analysis (GSEA) of genes associated with destabilized MT and cytoskeleton. **A.** The plot shows enrichment of genes associated with positive MT disassembly in PKCδ KO compared to Ramos WT cells. Vertical black lines indicate where genes overexpressed by MT depolymerization fall in the ranked list, (p-value 0.011 and FDR 0.81). **B.** As indication for the disconnection from the actin-cytoskeleton to the plasma membrane the plot shows the decrease of genes essential for the localization of the portion of the cytoskeleton that lies just beneath to the plasma membrane in PKCδ KO compared to Ramos WT cells, (p-value 0.00018 and FDR 0.091). The Normalized Enrichment Score (blue line) considers the ranked list of expression differences between two independent unstimulated PKCδ KO Ramos cell clones and Ramos WT cells.

**Supplementary figure 3:**
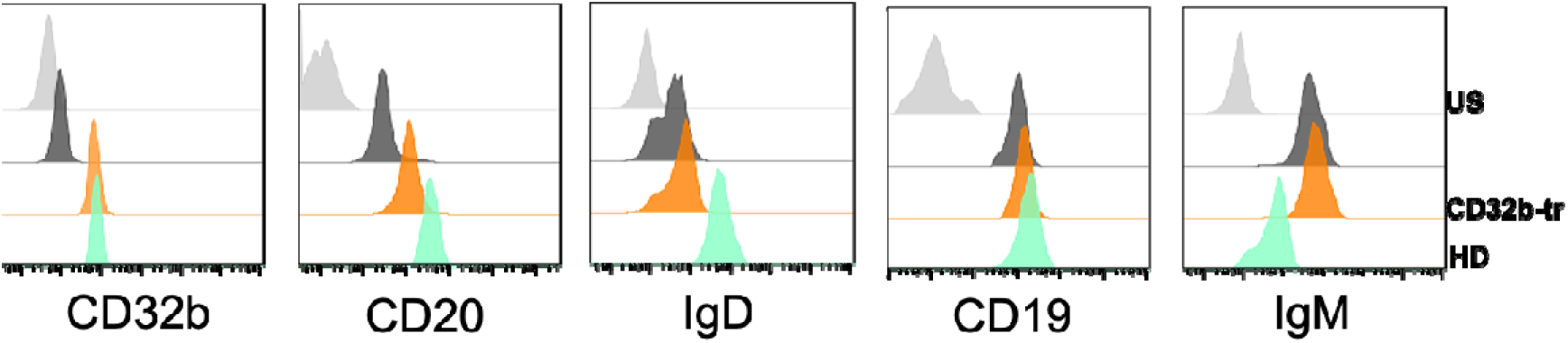
Due to the expression of the FcγRIIb receptor on Ramos, the surface abundance of critical B cell proteins more closely resembles that of naive B cells. Flow cytometry analysis shows the comparison of the expression of B cell surface protein abundance on healthy donor (HD) negative selected naïve B cells (green), Ramos (black) and FcγRIIb receptor transfected (CD32b-tr, orange) Ramos cells. Unstained (US) control in grey).

**Supplementary figure 4:**
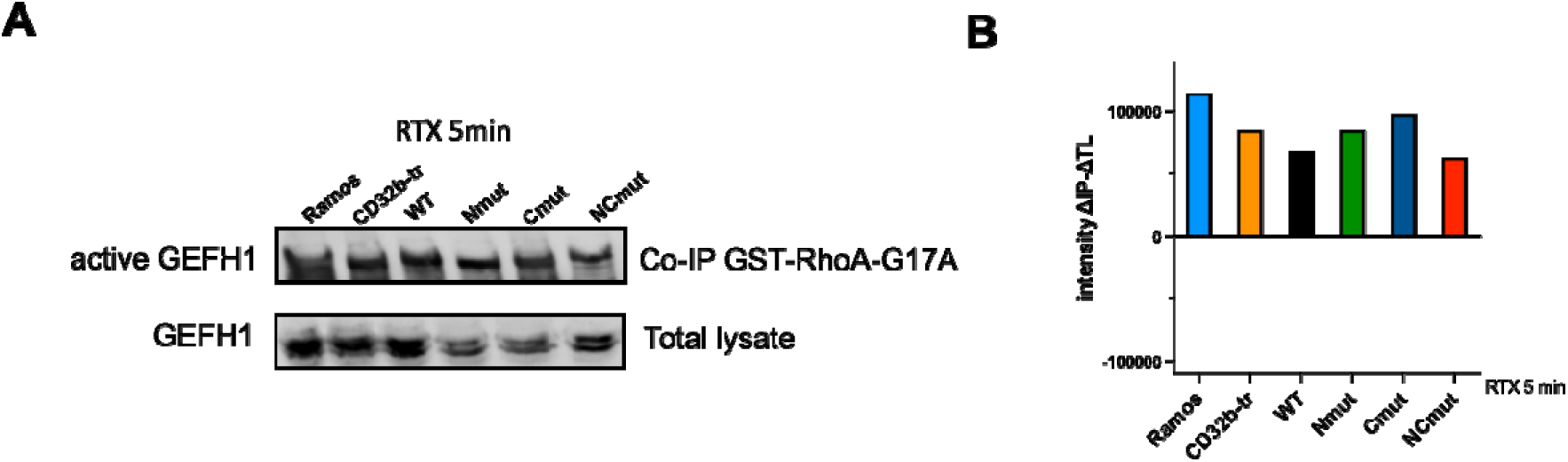
Western blot analysis of free GEF-H1 in the lysate of Ramos and CD32b-tr cells as well as reconstituted Ramos cells expressing CD20 WT, Nmut, Cmut or NCmut after activation with RTX for 5 min. The amount of GEF-H1 purified with the GST-RhoAG17A beads and in the total lysate is shown at the top and bottom, respectively. **F.** Quantification of the Western blot data with Image Studio Lite (Licor) showing the difference of signal intensity in the immunoprecipitate (IP) versus that of the total lysates (TL).

**Supplementary figure 5:**
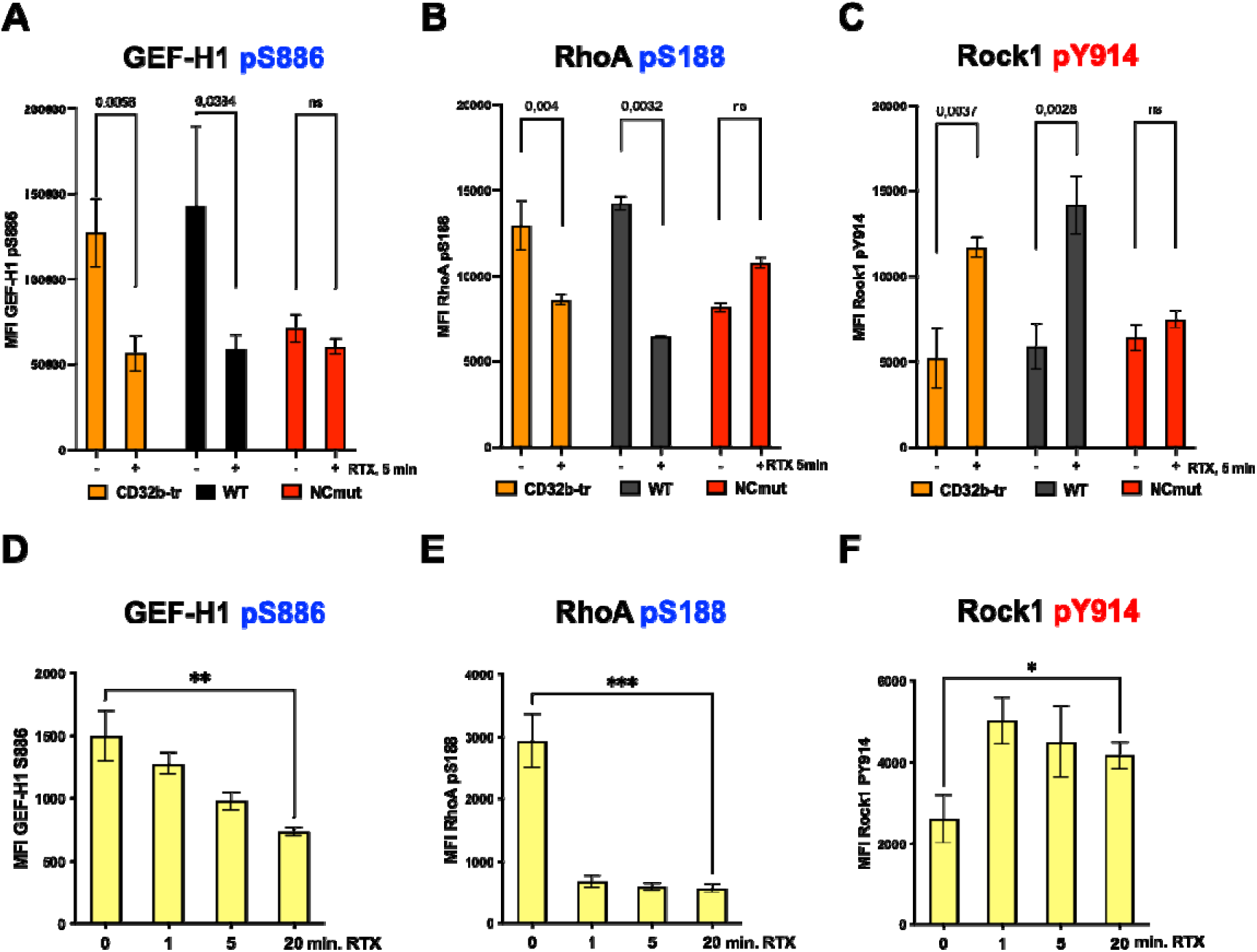
Intracellular phospho-flow cytometry analysis of intracellular signal proteins upon RTX treatment for 5 min compared to unstimulated CD20 WT and NCmut S>A transfectant show: **A** Loss of inhibitory GEF-H1 phosphorylation Serine^886^, which is a prerequisite for the activation of RhoA. **B** Loss of inhibitory RhoA Serine^188^ phosphorylation, which renders RhoA active. RhoA-GTP allows the increase of **C** active Rock1, phosphorylated at Y^914^. Students t-tests calculated with PRISM 10, show significant differences between the resting and stimulated state of CD32b-tr and WT cells, but not between S>A mutants. Representative NCmut CD20 mutant is presented. **D-F** Intracellular phospho-flow of the same protein residues as in **A-C** but on HD naive B cells and as a time course after RTX stimulation for 0, 1, 5, 20 min. **D** inhibitory GEF-H1 S^886^ phosphorylation, **E** inhibitory RhoA S^188^, and **F** activatory Rock1 Y914 phosphorylation. Significant differences between the groups were calculated with PRISM 10, one-way ANOVA. Activatory phosphorylations are depicted in red, inhibitory are shown in blue.

**Supplementary figure 6:**
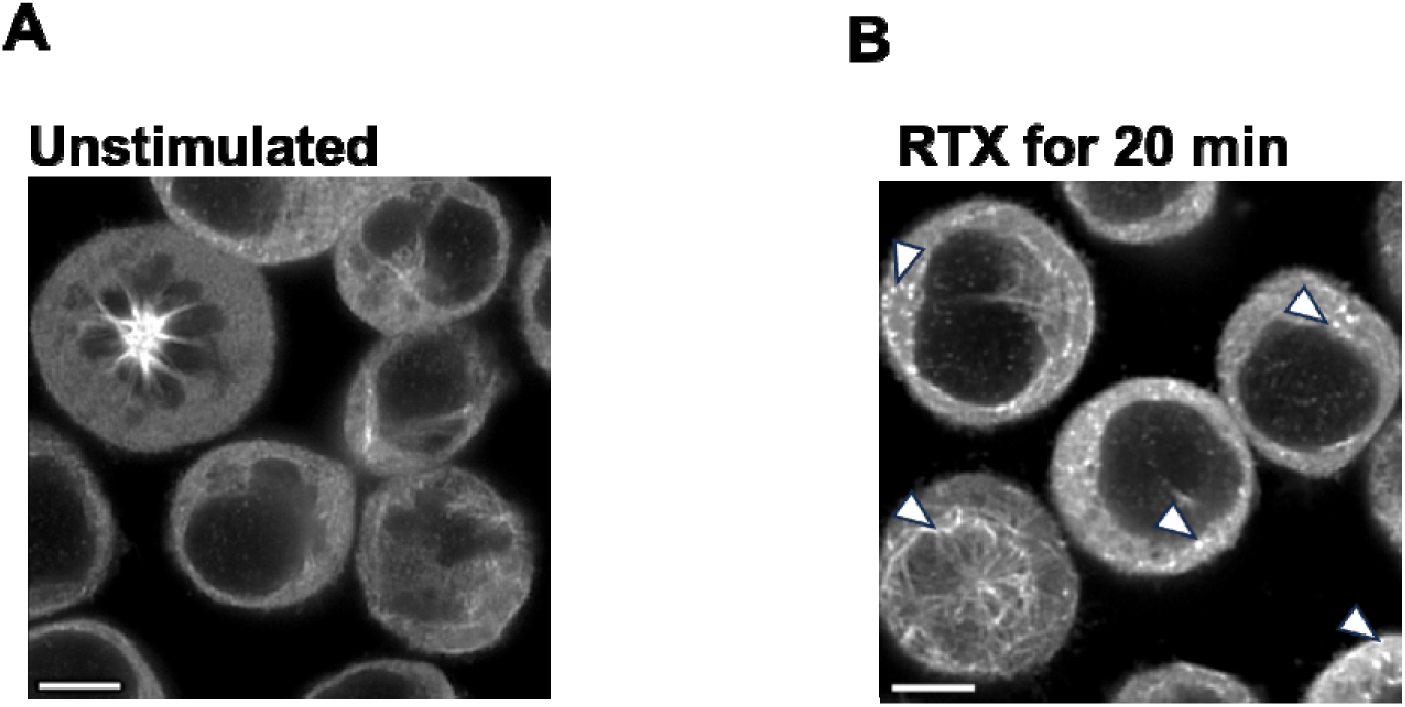
Airyscan confocal microscopy images of Ramos CD32b-tr cells **A.** unstimulated or **B.** incubated with RTX for 20 min. Cells were fixed and stained for α-tubulin with Alexa Fluor 555 to visualize MTs. Images present the mid-section of an Airyscan processed z-stack. White arrows in **B.** indicate the peeling of the curved protofilaments away from MT in the treated cells. Scale bar: 5µm.

**Supplementary figure 7:**
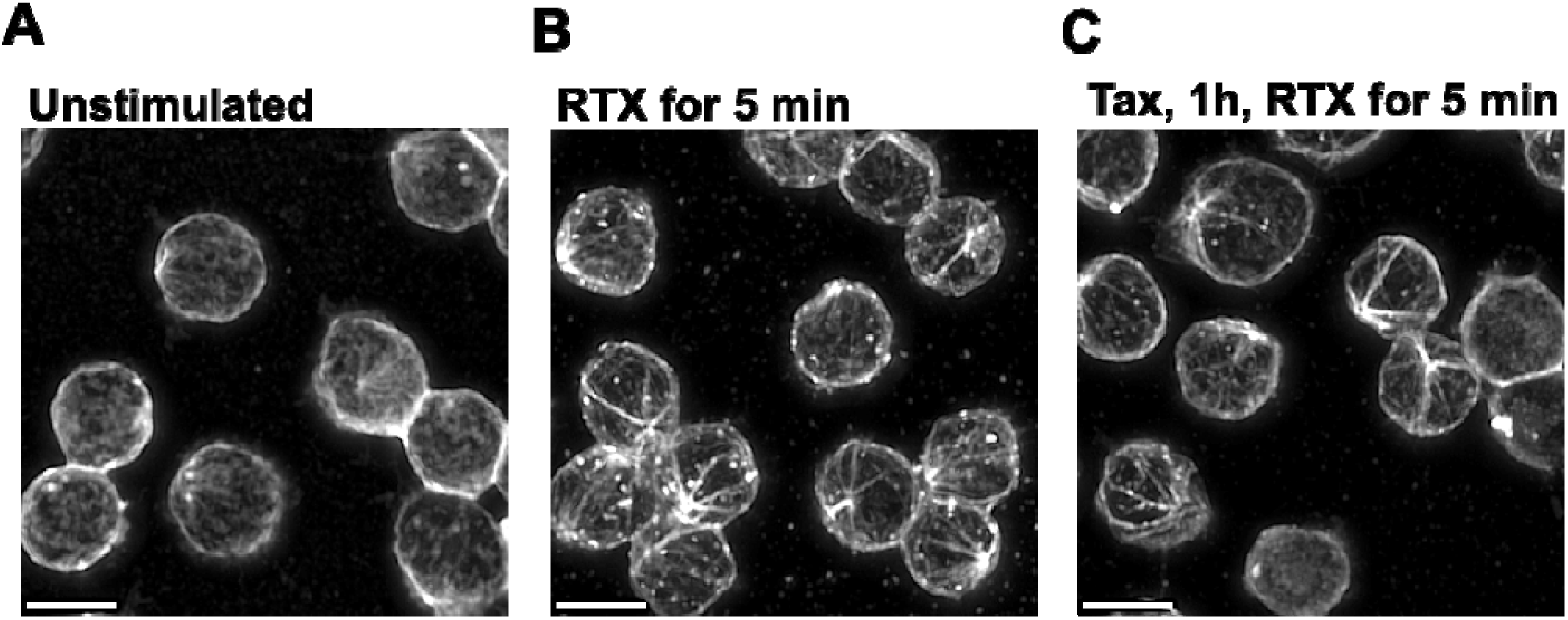
Maximum intensity projection of Airyscan (SR) confocal z-stacks of HD naïve B cells **A.** untreated, **B.** upon exposure to RTX for 5 min or **C**. with Taxol pretreatment for 1h before exposure to RTX for 5 min. Cells were fixed and stained for α-tubulin with Alexa Fluor 555. In **B.** background is brightened due to higher presence of dead or destroyed cells. Scale bar: 5µm.

## References

1. Pelanda R, Greaves SA, Alves da Costa T, Cedrone LM, Campbell ML, Torres RM. B-cell intrinsic and extrinsic signals that regulate central tolerance of mouse and human B cells*. Immunol Rev. 2022;307(1):12–26. doi:10.1111/imr.13062

2. Nemazee D. Mechanisms of central tolerance for B cells. Nat Rev Immunol. 2017;17(5):281–294. doi:10.1038/nri.2017.19

3. Kläsener K, Maity PC, Hobeika E, Yang J, Reth M. B cell activation involves nanoscale receptor reorganizations and inside-out signaling by Syk. Elife. 2014;3:e02069. doi:10.7554/eLife.02069

4. Depoil D, Fleire S, Treanor BL, et al. CD19 is essential for B cell activation by promoting B cell receptor-antigen microcluster formation in response to membrane-bound ligand. Nat Immunol. 2008;9(1):63–72. doi:10.1038/ni1547

5. Mattila PK, Feest C, Depoil D, et al. The Actin and Tetraspanin Networks Organize Receptor Nanoclusters to Regulate B Cell Receptor-Mediated Signaling. Immunity. 2013;38(3):461–474. doi:10.1016/j.immuni.2012.11.019

6. Horikawa K, Nishizumi H, Umemori H, Aizawa S, Takatsu K, Yamamoto T. Distinctive Roles of Fyn and Lyn in IgD-and IgM-Mediated Signaling. Vol 11.; 1999.

7. Becker M, Hobeika E, Jumaa H, Reth M, Maity PC. CXCR4 signaling and function require the expression of the IgD-class B-cell antigen receptor. Proceedings of the National Academy of Sciences. Published online May 1, 2017. doi:10.1073/PNAS.1621512114

8. Geisberger R, Lamers M, Achatz G. The riddle of the dual expression of IgM and IgD. Immunology. 2006;118(4):429–437. doi:10.1111/j.1365-2567.2006.02386.x

9. Maity PC, Blount A, Jumaa H, Ronneberger O, Lillemeier BF, Reth M. B cell antigen receptors of the IgM and IgD classes are clustered in different protein islands that are altered during B cell activation. Sci Signal. 2015;8(394):ra93. doi:10.1126/scisignal.2005887

10. Rougé L, Chiang N, Steffek M, et al. Structure of CD20 in complex with the therapeutic monoclonal antibody rituximab. Science (1979). 2020;367(6483):1224–1230. doi:10.1126/science.aba3526

11. Deans JP, Li H, Polyak MJ. CD20-mediated apoptosis: signalling through lipid rafts. Immunology. 2002;107(2):176–182. Accessed July 16, 2015. http://www.pubmedcentral.nih.gov/articlerender.fcgi?artid=1782791&tool=pmcentrez&rendertype=abstract

12. Reinhardt SCM, Masullo LA, Baudrexel I, et al. Ångström-resolution fluorescence microscopy. Nature. 2023;617(7962):711–716. doi:10.1038/s41586-023-05925-9

13. Einfeld DA, Brown JP, Valentine MA, Clark EA, Ledbetter JA. Molecular cloning of the human B cell CD20 receptor predicts a hydrophobic protein with multiple transmembrane domains. EMBO J. 1988;7(3):711–717. doi:10.1002/j.1460-2075.1988.tb02867.x

14. Pavlasova G, Mraz M. The regulation and function of CD20: An “enigma” of B-cell biology and targeted therapy. Haematologica. 2020;105(6):1494–1506. doi:10.3324/haematol.2019.243543

15. Riley JK, Sliwkowski MX. CD20: a gene in search of a function. Semin Oncol. 2000;27(6 Suppl 12):17–24.

16. Müller M, Gräbnitz F, Barandun N, et al. Light-mediated discovery of surfaceome nanoscale organization and intercellular receptor interaction networks. Nat Commun. 2021;12(1). doi:10.1038/s41467-021-27280-x

17. Arp AB, Abel Gutierrez A, ter Beest M, et al. CD70 recruitment to the immunological synapse is dependent on CD20 in B cells. Proceedings of the National Academy of Sciences. 2025;122(16). doi:10.1073/pnas.2414002122

18. Kosmas C, Stamatopoulos K, Stavroyianni N, Tsavaris N, Papadaki T. Anti-CD20-based therapy of B cell lymphoma: State of the art. Leukemia. 2002;16(10):2004–2015. doi:10.1038/sj.leu.2402639

19. Smith MR. Rituximab (monoclonal anti-CD20 antibody): Mechanisms of action and resistance. Oncogene. 2003;22(47):7359–7368. doi:10.1038/sj.onc.1206939

20. Margoni M, Preziosa P, Filippi M, Rocca MA. Anti-CD20 therapies for multiple sclerosis: current status and future perspectives. J Neurol. 2022;269(3):1316–1334. doi:10.1007/s00415-021-10744-x

21. Jazirehi AR, Bonavida B. Cellular and molecular signal transduction pathways modulated by rituximab (rituxan, anti-CD20 mAb) in non-Hodgkin’s lymphoma: Implications in chemosensitization and therapeutic intervention. Oncogene. 2005;24(13):2121–2143. doi:10.1038/sj.onc.1208349

22. Klein C, Jamois C, Nielsen T. Anti-CD20 treatment for B-cell malignancies: current status and future directions. Expert Opin Biol Ther. 2021;21(2):161–181. doi:10.1080/14712598.2020.1822318

23. Roubaud-Baudron C, Pagnoux C, Méaux-Ruault N, et al. Rituximab maintenance therapy for granulomatosis with polyangiitis and microscopic polyangiitis. Journal of Rheumatology. 2012;39(1):125–130. doi:10.3899/jrheum.110143

24. Payandeh Z, Bahrami AA, Hoseinpoor R, et al. The applications of anti-CD20 antibodies to treat various B cells disorders. Biomedicine and Pharmacotherapy. 2019;109:2415–2426. doi:10.1016/j.biopha.2018.11.121

25. Boross P, Leusen JHW. Mechanisms of action of CD20 antibodies. Am J Cancer Res. 2012;2(6):676–690.

26. Kläsener K, Jellusova J, Andrieux G, et al. CD20 as a gatekeeper of the resting state of human B cells. Proc Natl Acad Sci U S A. 2021;118(7). doi:10.1073/pnas.2021342118

27. Salzer E, Santos-Valente E, Keller B, Warnatz K, Boztug K. Protein Kinase C δ: a Gatekeeper of Immune Homeostasis. J Clin Immunol. 2016;36(7):631–640. doi:10.1007/s10875-016-0323-0

28. Lim PS, Sutton CR, Rao S. Protein kinase C in the immune system: From signalling to chromatin regulation. Immunology. 2015;146(4):508–522. doi:10.1111/imm.12510

29. Pracht C, Minguet S, Leitges M, Reth M, Huber M. Association of protein kinase C-δ with the B cell antigen receptor complex. Cell Signal. 2007;19(4):715–722. doi:10.1016/j.cellsig.2006.07.023

30. Sjef Verbeek J, Hirose S, Nishimura H. The complex association of fcγriib with autoimmune susceptibility. Front Immunol. 2019;10(OCT). doi:10.3389/fimmu.2019.02061

31. Karnell JL, Dimasi N, Karnell FG, et al. CD19 and CD32b Differentially Regulate Human B Cell Responsiveness. The Journal of Immunology. 2014;192(4):1480–1490. doi:10.4049/jimmunol.1301361

32. McCaleb MR, Miranda AM, Ratliff KC, Torres RM, Pelanda R. CD19 Is Internalized Together with IgM in Proportion to B Cell Receptor Stimulation and Is Modulated by Phosphatidylinositol 3-Kinase in Bone Marrow Immature B Cells. Immunohorizons. 2023;7(1):49–63. doi:10.4049/immunohorizons.2200092

33. Kläsener K, Yang J, Reth M. Study B Cell Antigen Receptor Nano-Scale Organization by in Situ Fab Proximity Ligation Assay. Vol 1707.; 2018. doi:10.1007/978-1-4939-7474-0_12

34. Tinti M, Johnson C, Toth R, Ferrier DEK, Mackintosh C. Evolution of signal multiplexing by 14-3-3-binding 2R-ohnologue protein families in the vertebrates. Open Biol. 2012;2(7):120103. doi:10.1098/rsob.120103

35. Pitasse-Santos P, Hewitt-Richards I, Abeywickrama Wijewardana Sooriyaarachchi MD, Doveston RG. Harnessing the 14-3-3 protein–protein interaction network. Curr Opin Struct Biol. 2024;86. doi:10.1016/j.sbi.2024.102822

36. Omasits U, Ahrens CH, Müller S, Wollscheid B. Protter: interactive protein feature visualization and integration with experimental proteomic data. Bioinformatics. 2014;30(6):884–886. doi:10.1093/bioinformatics/btt607

37. Ivaldi C, Martin BR, Kieffer-Jaquinod S, et al. Proteomic analysis of S-Acylated proteins in human B cells reveals palmitoylation of the immune regulators CD20 and CD23. PLoS One. 2012;7(5). doi:10.1371/journal.pone.0037187

38. Li Z, Ju X, Silveira PA, Abadir E, Hsu W hsun. CD83[: Activation Marker for Antigen Presenting Cells and Its Therapeutic Potential. 2019;10(June):1–9. doi:10.3389/fimmu.2019.01312

39. Ren Y, Li R, Zheng Y, Busch H. Cloning and Characterization of GEF-H1, a Microtubule-Associated Guanine Nucleotide Exchange Factor for Rac and Rho GTPases*. Vol 273.; 1998. http://www.jbc.org

40. Zenke FT, Krendel M, DerMardirossian C, King CC, Bohl BP, Bokoch GM. p21-activated Kinase 1 Phosphorylates and Regulates 14-3-3 Binding to GEF-H1, a Microtubule-localized Rho Exchange Factor. Journal of Biological Chemistry. 2004;279(18):18392–18400. doi:10.1074/jbc.M400084200

41. Krendel M, Zenke FT, Bokoch GM. Nucleotide exchange factor GEF-H1 mediates cross-talk between microtubules and the actin cytoskeleton. Nat Cell Biol. 2002;4(4):294–301. doi:10.1038/ncb773

42. Meiri D, Marshall CB, Mokady D, et al. Mechanistic insight into GPCR-mediated activation of the microtubule-associated RhoA exchange factor GEF-H1. Nat Commun. 2014;5. doi:10.1038/ncomms5857

43. Kashyap AS, Fernandez-Rodriguez L, Zhao Y, et al. GEF-H1 Signaling upon Microtubule Destabilization Is Required for Dendritic Cell Activation and Specific Anti-tumor Responses. Cell Rep. 2019;28(13):3367–3380.e8. doi:10.1016/j.celrep.2019.08.057

44. Kopf A, Kiermaier E. Dynamic Microtubule Arrays in Leukocytes and Their Role in Cell Migration and Immune Synapse Formation. Front Cell Dev Biol. 2021;9. doi:10.3389/fcell.2021.635511

45. Kosoff R, Chows HY, Radus M, Chernoffs J. Pak2 kinase restrains mast cell FcεRI receptor signaling through modulation of rho protein guanine nucleotide exchange factor (GEF) activity. Journal of Biological Chemistry. 2013;288(2):974–983. doi:10.1074/jbc.M112.422295

46. Waheed F, Speight P, Dan Q, Garcia-Mata R, Szaszi K. Affinity precipitation of active Rho-GEFs using a GST-tagged mutant Rho protein (GST-RhoA(G17A)) from epithelial cell lysates. J Vis Exp. 2012;(61). doi:10.3791/3932

47. Sluchanko NN, Gusev NB. 14-3-3 Proteins and regulation of cytoskeleton. Biochemistry (Moscow). 2010;75(13):1528–1546. doi:10.1134/S0006297910130031

48. Brandwein D, Wang Z. Interaction between Rho GTPases and 14-3-3 proteins. Int J Mol Sci. 2017;18(10). doi:10.3390/ijms18102148

49. Julian L. Investigating the Role of Caspase Cleavage of ROCK1 in Tissue Homeostasis and Tumour Development.; 2015.

50. Olson MF. Rho-associated coiled-coil containing kinases (ROCK): structure, regulation, and functions. Small GTPases. 2014;5:e29846. doi:10.4161/sgtp.29846

51. Birkenfeld J, Nalbant P, Yoon SH, Bokoch GM. Cellular functions of GEF-H1, a microtubule-regulated Rho-GEF: is altered GEF-H1 activity a crucial determinant of disease pathogenesis? Trends Cell Biol. 2008;18(5):210–219. doi:10.1016/j.tcb.2008.02.006

52. Pineau J, Pinon L, Mesdjian O, Fattaccioli J, Duménil AML, Pierobon P. Microtubules restrict F-actin polymerization to the immune synapse via GEF-H1 to maintain polarity in lymphocytes. Published online January 3, 2022. doi:10.1101/2022.01.03.473915

53. Mandelkow EM, Mandelkow E, Milligan RA. Microtubule Dynamics and Microtubule Caps: A Time-Resolved Cryo-Electron Microscopy Study.

54. Downing KH, Nogales E. Cryoelectron Microscopy Applications in the Study of Tubulin Structure, Microtubule Architecture, Dynamics and Assemblies, and Interaction of Microtubules with Motors. In: Methods in Enzymology. Vol 483. Academic Press Inc.; 2010:121–142. doi:10.1016/S0076-6879(10)83006-X

55. Basu A, Worth F, States U. Two Faces of Protein Kinase C δ[: The Contrasting Roles of PKC δ in Cell Survival and Cell Death. 2015;(August). doi:10.1100/tsw.2010.214

56. Gada KD, Logothetis DE. PKC regulation of ion channels: The involvement of PIP2. Journal of Biological Chemistry. 2022;298(6). doi:10.1016/j.jbc.2022.102035

57. Miyamoto. Increased proliferation of B cells and auto-immunity in mice lacking protein kinase Cd. Nature. 2002;(volume 416):865.

58. Sun Kuehn H, Niemela JE, Rangel-Santos A, et al. Loss-of-function of the protein kinase C d (PKCd) causes a B-cell lymphoproliferative syndrome in humans. Blood. 2013;121(16):3117–3125. doi:10.1182/blood

59. Chari R, Kim S, Murugappan S, Sanjay A, Daniel JL, Kunapuli SP. Lyn, PKC-δ, SHIP-1 interactions regulate GPVI-mediated platelet-dense granule secretion. Blood. 2009;114(14):3056–3063. doi:10.1182/blood-2008-11-188516

60. Kajimoto T, Sawamura S, Tohyama Y, Mori Y, Newton AC. Protein kinase C δ-specific activity reporter reveals agonist-evoked nuclear activity controlled by Src family of kinases. Journal of Biological Chemistry. 2010;285(53):41896–41910. doi:10.1074/jbc.M110.184028

61. Johnson C, Crowther S, Stafford MJ, Campbell DG, Toth R, MacKintosh C. Bioinformatic and experimental survey of 14-3-3-binding sites. Biochemical Journal. 2010;427(1):69–78. doi:10.1042/BJ20091834

62. Pennington K, Chan T, Torres M, Andersen J. The dynamic and stress-adaptive signaling hub of 14-3-3: emerging mechanisms of regulation and context-dependent protein–protein interactions. Oncogene. 2018;37(42):5587–5604. doi:10.1038/s41388-018-0348-3

63. Paul AL, Denison FC, Schultz ER, Zupanska AK, Ferl RJ. 14-3-3 Phosphoprotein interaction networks -Does isoform diversity present functional interaction specification? Front Plant Sci. 2012;3(AUG). doi:10.3389/fpls.2012.00190

64. Joo E, Olson MF. Regulation and functions of the RhoA regulatory guanine nucleotide exchange factor GEF-H1. Small GTPases. 2021;12(5-6):358–371. doi:10.1080/21541248.2020.1840889

65. Azoitei ML, Noh J, Marston DJ, et al. Spatiotemporal dynamics of GEF-H1 activation controlled by microtubule- And Src-mediated pathways. Journal of Cell Biology. 2019;218(9):3077–3097. doi:10.1083/JCB.201812073

66. Mok YK, Lo KWH, Zhang M. Structure of Tctex-1 and Its Interaction with Cytoplasmic Dynein Intermediate Chain. Journal of Biological Chemistry. 2001;276(17):14067–14074. doi:10.1074/jbc.M011358200

67. Meiri D, Marshall CB, Greeve MA, et al. Mechanistic Insight into the Microtubule and Actin Cytoskeleton Coupling through Dynein-Dependent RhoGEF Inhibition. Mol Cell. 2012;45(5):642–655. doi:10.1016/j.molcel.2012.01.027

68. van de Ven AAJM, Compeer EB, Bloem AC, et al. Defective calcium signaling and disrupted CD20-B-cell receptor dissociation in patients with common variable immunodeficiency disorders. J Allergy Clin Immunol. 2012;129(3):755–761.e7. doi:10.1016/j.jaci.2011.10.020

69. Li H, Ayer LM, Polyak MJ, et al. The CD20 calcium channel is localized to microvilli and constitutively associated with membrane rafts: Antibody binding increases the affinity of the association through an epitope-dependent cross-linking-independent mechanism. Journal of Biological Chemistry. 2004;279(19):19893–19901. doi:10.1074/jbc.M400525200

70. Schober R, Waldherr L, Schmidt T, et al. STIM1 and Orai1 regulate Ca 2+ microdomains for activation of transcription. Biochim Biophys Acta Mol Cell Res. 2019;1866(7):1079–1091. doi:10.1016/j.bbamcr.2018.11.001

71. Fais S, Malorni W. Leukocyte uropod formation and membrane/cytoskeleton linkage in immune interactions. J Leukoc Biol. 2003;73(5):556–563. doi:10.1189/jlb.1102568

72. Saltukoglu D, Özdemir B, Holtmannspötter M, et al. Plasma membrane topography governs the 3D dynamic localization of IgM B cell antigen receptor clusters. EMBO J. 2023;42(4):e112030. doi:10.15252/embj.2022112030

73. Mattila PK, Batista FD, Treanor B. Dynamics of the actin cytoskeleton mediates receptor cross talk: An emerging concept in tuning receptor signaling. J Cell Biol. 2016;212(3).

74. Wang J, Lin F, Wan Z, et al. I M M U N O L O G Y Profiling the Origin, Dynamics, and Function of Traction Force in B Cell Activation.; 2018. https://www.science.org

75. Sanchez-Madrid F. SJ. Bringing up the rear- defining the roles of the uropod. Nat Rev Mol Cell Biol. 2009;Volume 10:353–359.

76. Roberts PJ, Mitin N, Keller PJ, et al. Rho family GTPase modification and dependence on CAAX motif-signaled posttranslational modification. Journal of Biological Chemistry. 2008;283(37):25150–25163. doi:10.1074/jbc.M800882200

77. Lehtimäki JI, Rajakylä EK, Tojkander S, Lappalainen P. Generation of stress fibers through myosin-driven reorganization of the actin cortex. Elife. 2021;10:1–43. doi:10.7554/eLife.60710

78. Rey M, Sanchez-Madrid F, Valenzuela-Fernandez A. The Role of Actomyosin and the Microtubular Network in Both the Immunological Synapse and T Cell Activation. Vol 12.; 2007.

79. Reth M. Discovering immunoreceptor coupling and organization motifs. Front Immunol. 2023;14. doi:10.3389/fimmu.2023.1253412

80. Shanehbandi D, Majidi J, Kazemi T, Baradaran B, Aghebati-Maleki L. CD20-based Immunotherapy of B-cell Derived Hematologic Malignancies. Curr Cancer Drug Targets. 2017;17(5):423–444. doi:10.2174/1568009617666170109151128

81. Deans JP, Li H, Polyak MJ. CD20-Mediated Apoptosis: Signalling through Lipid Rafts.; 2002.

82. Chang YC, Nalbant P, Birkenfeld J, Chang ZF, Bokoch GM. 3GEF-H1 couples nocodazole-induced microtubule disassembly to cell contractility via RhoA. Mol Biol Cell. 2008;19(5):2147–2153. doi:10.1091/mbc.E07-12-1269

83. Kozlova V, Ledererova A, Ladungova A, et al. CD20 is dispensable for B-cell receptor signaling but is required for proper actin polymerization, adhesion and migration of malignant B cells. PLoS One. 2020;15(3). doi:10.1371/journal.pone.0229170

84. Ghosh A, Meub M, Helmerich DA, et al. Decoding the molecular interplay of CD20 and therapeutic antibodies with fast volumetric nanoscopy. Science (1979). 2025;387(6730). doi:10.1126/science.adq4510

85. Hiraga J, Tomita A, Sugimoto T, et al. Down-regulation of CD20 expression in B-cell lymphoma cells after treatment with rituximab-containing combination chemotherapies: its prevalence and clinical significance. Published online 2009. doi:10.1182/blood-2008-08

86. Marshalek JP, Dragan M, Tomassetti S, Labarbera K, Amaya A, Har ; E19537 Publication Only CD20 and CD19 Expression Loss in Relapsed or Refractory B-Cell Non-Hodgkin Lymphoma: A Retrospective Cohort.; 2022.

87. Däbritz JHM, Yu Y, Milanovic M, et al. CD20-targeting immunotherapy promotes cellular senescence in B-cell lymphoma. Mol Cancer Ther. 2016;15(5):1074–1081. doi:10.1158/1535-7163.MCT-15-0627

