## Extended Table1 for "Microtubule anchoring and coupling of CD20 to the RhoA/Rock1 pathway"

**Extended Table 1**

CD20-WT geneblock
5'/5Phos/TCTACCCAATACTGTTACAGCATACAATCTCTGTTCTTGGGCATTTTGTCAGTGA
TGCTGATCTTTGCCTTCTTCCAGGAACTTGTAATAGCTGGCATCGTTGAGAATGAATGGA
AAATAACGTGCTCCAGACCCTTTGCCTTCTTCCAGGAACTTGTAATAGCTGGCATCGTTG
AGAATGAATGGAAAATAACGTGCTCCAGACCCAAATCTAACATAGTTCTCCTGTCAGCAG
AAGAAAAAAAAGAACAGACTATTGAAATAAAAGAAGAAGTGGTTAAATCTAACATAGTTCT
CCTGTCAGCAGAAGAAAAAAAAGAACAGACTATTGAAATAAAAGAAGAAGTGGTTGGGC
TAACTGAAACATCTTCCCAACCAAAGAATGAAGAAGACATTGAAATTATTCCAATCCAAGA
AGAGGAAGGGCTAACTGAAACATCTTCCCAACCAAAGAATGAAGAAGACATTGAAATTAT
TCCAATCCAAGAAGAGGAAGAAGAAGAAACAGAGACGAACTTTCCAGAACCTCCCCAA
GATCAGGAATCCTCACCAATAGAAAATGACAGCGAAGAAGAAACAGAGACGAACTTTCC
AGAACCTCCCCAAGATCAGGAATCCTCACCAATAGAAAATGACAGCTCTCCTGATTACAA
GGATGACGACGATAAGTGACGCCCCCCCCCCCTAACGTTACTGGCCGAAGCCGCTTGG
TCTCCTGATTACAAGGATGACGACGATAAGTGACGCCCCCCCCCCCTAACGTTACTGGC
CGAAGCCGCTTGGAATAAGGCCGGTGTGCGTTTGTCTATATGTTATTTTCCACCATATTG
CCGTCTTTTGGCAATAAGGCCGGTGTGCGTTTGTCTATATGTTATTTTCCACCATATTGCC
GTCTTTTGGC-3'

CD20-C-term Cmut geneblock
5'/5Phos/GTTGAGAATGAATGGAAAAGAACGTGCGCCAGACCCAAAGCTAACATAGTTCT
CCTGTCAGCAGAAGAAAAAAAAGAACAGACTATTGAAATAAAAGAAGAAGTGGTTGGGC
TAACTGAAACATCTTCCCAACCAAAGAATGAAGAAGACATTGAAATTATTCCAATCCAAGA
AGAGGAAGAAGAAGAAACAGAGACGAACTTTCCAGAACCTCCCCAAGATCAGGAATCCT
CACCAATAGAAAATGACAGCTCTCCT-3'

CD20-N-term Nmut geneblock
5'/5Phos/TCCTCCATCCGCCCCGTCTCTCCCCCTTGAACCTCCTCGTTCGACCCCGCCTC
GATCCTCCCTTTATCCAGCCCTCACTCCTTCTCTAGGCGCCGGAATTAGATCTCTCGAGG
TTAACGAATTCATGACAACACCCAGAAATGCAGTAAATGGGACTTTCCCGGCAGAGCCA
ATGAAAGGCCCTATTGCTATGCAATCTGGTCCAAAACCACTCTTCAGGAGGATGGCTGC
ACTGGTGGGCCCCACGCAAAGCTTCTTCATGAGGGAAGCTAAGACTTTGGGGGCTGTC
CAGATTATGAATGGGCTCTTCCACATTGCCCTGGGGGGTGTTGTGATGATCCCAGCAGG
GATCTATGCACCCATCTGTGTGACTGTGTGGTACCCTCTCTGGGGAGGC-3’
