## Supplementary material for "Microtubule anchoring and coupling of CD20 to the RhoA/Rock1 pathway": Materials and Methods

**Materials & Methods**

**Human naïve B cells**

Primary naïve B cells were obtained from fresh buffy coats, provided by the Institute for Transfusion Medicine and Gene Therapy, ITG, Freiburg. Peripheral blood mononuclear cells (PBMCs) were separated by Ficoll gradient centrifugation of whole blood and negatively selected using EasySep Human Naïve B Cell Isolation Kit (Stemcell). Prior to experiments, primary B cells were controlled for purity and rested overnight. The study was approved by the Institutional Review Board of the University Freiburg (Ethical votes No 507/16 and 336/16).

**Cell culture**

The human Burkitt lymphoma B cell line Ramos was obtained from American Type Culture Collection (ATCC, Ramos cat. CRL-1923, RRID: CVCL 1646) and stably transfected with ecotropic receptor (EcoR) to allow murine retroviral transfection with MMLV particles. Ramos B cells were cultured in RPMI 1640-Glutamaxx medium, 10% FCS (PAN, Biotech), 10 units/mL penicillin/streptomycin (Gibco), 20 mM hepes (Gibco), and 50 mM β-mercapto-ethanol (Sigma) in a humidified saturated atmosphere at 37 °C with 5% CO_2_.

For the quantitative phosphoproteomic study, Ramos B cells were labelled using stable isotype labelling by amino acids in cell culture (SILAC). Light labelling was performed using 12C6L-arginine/12C6-L-lysine. For medium-heavy and heavy labelling 13C6L-arginine/D4-L-lysine and 13C615N4-L-arginine/13C615N2-L-lysine were used, respectively.

**Generation of the Ramos CD32b (*FCGR2B*) KO and CD20 (*MS4A1*) KO cell lines**

The human genes for CD32b (*FCGR2B*) and CD20 (*MS4A1*) were rendered defective in the Ramos B cell system using the CRISPR/Cas9 system. Synthetic single guide RNAs (sgRNA)s targeting defined 20-bp sequences within the corresponding coding regions (agcacgttgatccactgggg for CD32b KO located in exon 4 of *FCGR2B* and gccctattgctatgcaatctgg within the exon 3 of MS4A1 for CD20 KO, respectively) were purchased form Integrated DNA Technology (IDT). The CRIPR-Cas components were delivered to the Ramos B cell line in form of ribonucleoprotein (RNP) complex, using the above mentioned sgRNAs pre-assembled with Cas9 protein (IDT) via the NEON Transfection System (Invitrogen) following the manufacturer’s instructions. In brief: Ramos cell medium was changed the day before transfection. 1 × 10^6^ Ramos cells were pelleted by centrifugation and resuspended in 9 μL buffer R (Neon), 1µL of RNP complex, and 2µL of electroporation enhancer (IDT). RNP complex contained diluted sgRNA (annealed crRNA and tracrRNA in a 1:1 ratio in IDTE buffer) and Cas9 endonuclease in buffer R (NEON). Electroporation was performed in 10 µL NEON tips at 1350 V, 30 ms, single pulse. The transfected cells were first recovered for 72hs at 37°C and 5% CO2 without antibiotics and then subjected to complete RPMI medium. Successful transfected Ramos cells were batch sorted using BioRad cell sorter in three different rounds and monitored throughout experiments. Inactivation of the target genes was verified by flow cytometry.

**Generation of the Ramos PKCδ (*PRKCD*) gene KO**

The sequences AACCCAATCATAGCAGAGC or GCTGAGTTCAGTGAGTGC, each targeting the *PRKCD* coding region were inserted into a lentivirus CRISPR vector (Addgene) to generate the corresponding sgRNA cassettes. The constructs were co-transfected with an mCherry expression vector into Ramos cells. The cells were incubated at 37°C for 24 hours in a humidified CO2 incubator. The following day, single cells were sorted for mCherry reporter gene expression with the BD FACSAria™ III Cell Sorter. The sorted single cells were cultured for 3-4 weeks in complete RPMI medium supplied with 10% FBS in a humidified CO_2_ incubator at 37°C. The expected *PRKCD* gene deficiency was verified by Sanger sequencing and the loss of PKCδ protein expression by Western blotting.

**Lentiviral transfection for the generation of CD32b-tr Ramos B cells**

The human embryonic kidney HEK 293T cells were cultured in complete DMEM GlutaMAX medium supplemented with 10% FCS, 10 mM HEPES, 10 μM sodium pyruvate, 50 units/mL penicillin and 50 µg/mL streptomycin in a humidified saturated atmosphere at 37 °C with 7.5% CO_2_. Lentivirus particles were obtained by co-transfecting HEK 293T cells with pLenti-hCD32b-IRES-GFP, pCMVΔR8.74 and pMD2vsvG plasmids using Polyethyleneimine (PEI, Polysciences). Viral supernatant was harvested, sterile filtered and combined 24h and 48h post-transfection. Lentiviral particles were enriched by centrifugation (4 hs, 10.000 g, 8 °C) after placing them in a 1:5 ratio on a 10% sucrose layer. The lentiviral pellet was resuspended in DMEM Glutamaxx without supplements and stored at –80 °C. The viral titers were assessed by determining the multiplicity of infection (MOI). Briefly, 5x10^4^ HEK 293T cells per well were seeded in a p24 well plate in 1 mL medium. An aliquot of the concentrated virus was diluted 1:100 in medium. Various volumes (0, 1, 5, 10, 25, and 50 µL) of lentivirus dilution were added to the cells. After 48h of incubation the GFP expression was analyzed by flow cytometry and lentivirus titer was calculated: Transduction units/mL = (number of cells x % of GFP^+^ cells x dilution factor) / (mL of lentivirus dilution).

**Construction of WT or S/A mutated CD20-flag expression vectors**

The CD20-flag vector was modified by geneblocks (IDT) encoding the N-terminal or C-terminal tails with S/A exchanges at the chosen 4 positions, respectively. Geneblocks were solved in IDTE buffer according manufacturer’s protocol and subsequently inserted in the pMIG CD20-flag-IRES-GFP vector using infusion cloning method. Plasmid were sequenced (GATC, eurofins) and used to generate the CD20WT, CD20Nmut, CD20Cmut and CD20NCmut Ramos cell lines (see extended table 1, ET1).

**Retroviral transfection of CD20 mutants:**

Murine retrovirus-containing supernatants were obtained by transfecting Phoenix-eco cells with LipoJet (SignaGen) according to manufacturer’s protocol and different pMIG-CD20-IRES-GFP constructs, or empty GFP-control plasmid. After 48h viral supernatant was collected, sterile filtered and used directly or stored in -80°C. For transduction 4x10^5^ newly generated CD20 KO Ramos cells were resuspended in 1 mL of viral supernatant containing Polybrene (1 µg/mL) and spin infected by centrifugation (180 min, 300g, 37°C). The viral supernatant was removed and cells were cultured and sorted for GFP expression (BioRad cell sorter).

**RNA sequencing**

Ramos wt or PKCδKO Ramos were cultured with or without IgM (10ug/ml) stimulation for 4 hours and then RNAs were extracted by RNeasy plus kit (Qiagen cat# 74143). RNA libraries for RNAseq were prepared using the NEBNext® Ultra™ II Directional RNA Library Prep Kit for Illumina (NEB cat # E7760L) according to manufacturer’s instructions and samples were sequenced 84-bases long on the NextSeq® 550 with high output flowcell (NextSeq® 500/550 High Output Kit v2).

**GEO data**

Downstream analyses were conducted using R (version 4.4.0). Raw read counts were normalized for library size and composition using the trimmed mean of M-values (TMM) method. Differential gene expression analysis comparing the unstimulated Ramos and PKCδ KO cell conditions was performed with the edgeR package (version 4.2.2)^88^. Gene set enrichment analysis was carried out using the clusterProfiler package (version 4.12.6)^89^ with gene sets from the Molecular Signatures Database (MSigDB, version 7)^90^. For all statistical analyses, significance was defined as an adjusted p-value < 0.05.

(GEO accession GSE300605 on [www.ncbi.nlm.nih.gov/geo/query/acc.cgi?acc=GSE300605](http://www.ncbi.nlm.nih.gov/geo/query/acc.cgi?acc=GSE300605))

**Flow cytometry analysis and phospho-flow detection**

For surface staining, 1–20 x 10^5^ cells were Fc-blocked (Human TruStain, Biolegend), stained with fluorescent antibodies on ice, washed twice, and then their fluorescence measured with a FACS Attune NxT (Life technologies). For intracellular staining and phospho-flow analysis, the cells were activated for the requested time points, immediately fixed with 4% PFA, washed twice and permeabilized with 0.5% saponin (Quillaja bark, Sigma) containing protease and phosphatase inhibitors (HALT, Thermo Fisher) for 30 min. Samples were washed and stained overnight at 4°C. If necessary secondary antibody staining was performed, the background was controlled using only the secondary antibody. Data were exported in FCS‐3.0 format and analyzed with FlowJo software (TreeStar).

**Western Blotting**

Unstimulated or activated Ramos B cells were collected and immediately lysed in 2x Laemmli or RIPA buffer for 30 min on ice. Equal amounts of adjusted and cleared lysates (Qubit 3.0; life technologies) were subjected to SDS–PAGE on 10% mini precast gels (7bioscience). After transfer, the PVDF membrane (GE Healthcare Amersham) was blocked with 5% BSA in PBS and 0.1% Tween20. The membrane was first exposed to primary antibodies specific for the proteins under study (CD20, 14-3-3 or GEF-H1) and then washed. A horseradish (HRP) peroxidase-conjugated goat anti-rabbit or goat anti-mouse antibody was used as secondary antibody before detection with ECL chemiluminescent substrate (Bio-Rad).

**Affinity purification of CD20-flag and GEF-H1**

The anti-flag immunoprecipitation was performed according to the manufacturer's protocol (chromotek). In brief, 10^7^ Ramos cells were lysed for 30 min in ice-cold lysis buffer supplemented with a protease/phosphatase inhibitor cocktail (HALT, Thermo Fisher). Lysates were centrifuged for 10 min at 17000 g and diluted in 300µL dilution buffer. 50 µL of total lysate were kept for the input analysis. Diluted lysates were rotated overnight with anti-flag agarose beads. Beads were then sedimented and washed extensively. Bound proteins were eluted with 2x SDS-PAGE Laemmli buffer and samples were boiled at 95°C for 5 min and subjected to SDS-PAGE and Western blotting.

RhoA(G17A), the nucleotide-binding deficient mutant of RhoA binds strongly to GEF-H1 and can be used for GEF-H1 purification and quantification. For this a glutathione S-transferase (GST) coupled RhoA(G17A) fusion protein was produced in E. coli bacteria and the GST-RhoA(G17A) loaded on Glutathione Sepharose beads. The GST-RhoA(G17A)-beads were mixed with Ramos B cell lysate and rotated overnight at 4°C. Beads were washed extensively with lysis buffer, boiled for 5 min and the cleared supernatant subjected to SDS-PAGE and western blotting.

**Proximity ligation assay (PLA)**

The Fab-PLA was performed as described earlier^33^. In brief: for Fab‐PLA, the F(ab)- fragments were prepared from the corresponding antibodies using the Pierce Fab Micro Preparation Kit (Thermo Fisher). The produced F(ab)-fragments or primary antibodies (1-PLA) coupled with PLA probemaker (Sigma‐Aldrich) after buffer exchange (Zeba™ spin desalting columns, Thermo Fisher). Ramos or HD naïve B cells were settled on polytetrafluoroethylene (PTFE) slides (Thermo Fisher Scientific) for 30 min at 37°C and kept unstimulated or activated with RTX [10µg / mL] for 5 min and then fixed for 20 min with 4% paraformaldehyde.

After extensive washings the slides were blocked for 30 min with blocking buffer (25 μg/mL sonicated salmon sperm DNA, 250 μg/mL BSA, 1 M glycine). PLA was performed with the Duolink In Situ Orange (Sigma-Aldrich). Slides were directly mounted with DAPI Fluoromount-G (Southern Biotech) to visualize the PLA signals in relation to the nuclei. For each experiment, a minimum of 1000 HD B cells or Ramos cells were analyzed with CellProfiler-3.0.0. In brief: metadata of images were exported and the intensity of the input frames were rescaled to the full intensity range in the RGB scheme. PLA signals (size of PLAdots 4-10 pixels) in a distance of 10 pixels to the Nuclei (size 15-80 pixel) with a threshold of 0.0075 of at least 1000 cells were measured and subjected to analysis.

**Imaging, Image analysis, and Data processing**

All microscope images for PLA were acquired using Leica DMi8 microscope equipped with an HC PL APO 63×/1.40-0.60 oil immersion objective lens and analyzed with CellProfiler 3.0.0. and Prism software (GraphPad, La Jolla, CA).

**Microtubule staining**

MT were stained as described^91^. In brief: Ramos B cells were settled on uncoated glass bottom chambers (IBIDI) for 30 min at 37°C, kept unstimulated or activated with RTX for 20 min and then fixed for 20 min with 4% paraformaldehyde and 0.1% glutaraldehyde. After blocking and permeabilization with 250 μg/mL BSA, 1 M glycine 0.5% saponine for 30 min cells were stained with anti-α tubulin (Sigma Aldrich) over night and with secondary anti mouse Alexa fluor-555 antibody (Invitrogen). Fixed Ramos cells in IBIDI chambers were kept in PBS until microscope imaging.

HD B cells were settled on PTFE slides (Thermo Fisher) for 30 min at 37°C, kept unstimulated, or activated with RTX [10µg/mL] for 5 min, with or without pretreatment with Taxfor 1h and then fixed for 20 min with 4% paraformaldehyde and 0.1% glutaraldehyde. After blocking and permeabilization with 250 μg/mL BSA, 1 M glycine 0.5% saponine for 30 min cells were stained with anti-α tubulin (Sigma Aldrich) over night and with secondary anti mouse Alexa fluor-555 antibody (Invitrogen). Slides were directly mounted with Fluoromount-G mounting-solution (Invitrogen) and subjected to microscope analysis.

**Microscopy**

The high-resolution visualization of MTs was performed using a LSM 880 Airyscan laser scanning confocal microscope attached to an inverted microscope Axio Observer Z1 (Carl Zeiss Microscopy). The Alexa Fluor 555 labelled α-tubulin was excited with 561nm, the emission was detected with the Airyscan detector in SuperResolution (SR) mode. For all cells z-stacks were acquired with a Plan-Apochromat 63x/1.40 Oil DIC M27 (Fig. S7) or a LD LCI Plan-Apochromat 40x/1.20 autocorr (Fig. S6) objective. The immersion medium was oil (63x) or water (40x).

Zeiss Software ZEN Black 2.3 SP1 FP3 (release 14.0.22.201) was used for acquisition. (acquisition settings for Fig. S7: EM filter SP615, pixel size xy:0.049µm, z-step size: 0.199µm, pixel dwell time 1µs, bi-directional scanning, for Fig. S6: no EM filter, pixel size xy:0.049µm, z-step size: 0.243µm or 1µm, pixel dwell time 1,92µs, bi-directional scanning). All images were processed with the above ZEN Black software using the Airyscan algorithm and with Zen Blue (Version 3.9.3) for subset and maximum intensity projection. Airyscan processed images shown-in Fig. S6 are single z-planes from an image z-stack and in Fig. S7 maximum intensity projections of the full image z-stack. The display gain is set to 0.75 and the intensity value to Best fit.

**Statistical analysis**

Students t-test was used for the experiments if not otherwise described to determine statistical significance provided by Prism10 software (GraphPad, La Jolla, CA).

**MT network stabilization or destabilization**

For the stabilization of MT HD naïve B cells or Ramos B cells were treated for 1 h with 100 nM docetaxel trihydrate (MedChemExpress) solved in DMSO and diluted in RPMI culture medium. For the destabilization of MT cells were treated for 1 h with 10 μM nocodazole (Selleck-Chemicals) solved in DMSO and diluted in RPMI culture medium.

**Rituximab (RTX) treatment**

RTX was kindly provided by F. Hoffmann-La Roche-AG. For naïve HD B cells or Ramos cell treatment RTX was diluted to a final concentration of 10 µg/mL and experiments were performed at least three times.

**Phosphopeptide enrichment**

SILAC-labeled Ramos B-cells (wt and PKCδ KO) were starved by serum depletion (FCS) for 30 min. Following starvation, cells were harvested by centrifugation at 1,200 rpm for 5 min, washed twice with starvation medium. As a control, mock treatment was performed using PBS. After activation, cells were rapidly snap-frozen in liquid nitrogen to preserve their signaling status. Cell lysis was performed using GdmHCl buffer (6 M GdmHCl, 100 mM Tris-HCl pH 8.5, 10 mM TCEP, and 40 mM chloroacetamide). The cell pellets were resuspended in the lysis buffer, followed by sonication (two cycles of 30 seconds). Protein denaturation was achieved by incubating the lysates at 95°C for 5 min. To remove cellular debris, the lysates were centrifuged at 3,500 x g for 30 min at 4°C. Protein concentration was determined using a Bradford assay. To precipitate proteins, four volumes of ice-cold acetone were added to the lysate. The protein precipitates were collected by centrifugation and washed with ice-cold acetone to remove residual contaminants. Proteins were digested using a combined Lys-C / trypsin digestion. Phosphopeptides were enriched using the Easyphos workflow. The phosphopeptide-enriched eluates were desalted using C18 StageTips (3M Company), pooled, and stored at -80°C until further analysis.

**LC-MS/MS analysis**

Peptide mixtures, reconstituted in 0.1% TFA, were analyzed by nano HPLC-ESI-MS/MS using an UltiMate 3000 RSLCnano HPLC system (Thermo Fisher Scientific, Dreieich, Germany) online coupled to an QExactive Plus mass spectrometer (Thermo Fisher Scientific, Bremen, Germany). The RSLC system was equipped with C18 trap columns (nanoEase M/Z Symmetry C18 Trap; 20 mm length, 180 µm inner diameter, 5 µm particle size, 100 Å pore size, Waters Corporation, Milford, MA) and an analytical C18 reversed-phase nano LC column (nanoEase M/Z HSS C18 T3; 250 mm length, 75 mm inner diameter, 1.8 µm particle size, 100 Å pore size, Waters Corporation, Milford, MA). A binary solvent system consisting of 0.1% (v/v) formic acid (FA) as solvent A and 80% (v/v) acetonitrile (ACN)/0.1% (v/v) FA as solvent B was employed for peptide separation. Peptide mixtures were loaded, washed and preconcentrated on the pre-column for 5 min using solvent A and a flow rate of 10 μL/min. A gradient was then applied at a flow rate of 300 nL/min ranging from 4 to 39% B in 140 min, 39–54% B in 15 min, 54%-95% in 5 min and 3 min at 95% B. Eluted peptides were transferred to a fused silica emitter for electrospray ionization enabled by a Nanospray Flex ion source with DirectJunction adaptor (Thermo Fisher Scientific), applying a spray voltage of 1.6 kV and a capillary temperature of 250°C. MS/MS data were acquired in data-dependent mode using the following parameters: MS precursor scans at m/z 370–1700 with a resolution of 70,000 (at m/z 400); automatic gain control (AGC) of 3 × 10^6^ ions; a maximum injection time (IT) of 60 ms; a top 12 method for higher-energy collisional dissociation of multiply charged precursor ions with a normalized collision energy of 28%. MS/MS scans from 200 to 2000 m/z were recorded at a resolution of 35,000. The AGC for MS/MS scans was set to 1 × 10^5^ with a maximum IT of 120 ms and a dynamic exclusion time of 45 s. Data are available via ProteomeXchange with identifier PXD063667.

**MS data analysis**

MaxQuant (version 2.6.7.0) with its integrated Andromeda search engine^92,93^ was employed. MS/MS data were searched against the human proteome set with isoforms downloaded from UniProt (105300 entries). Protein identification was performed using MaxQuant default settings (including carbamidomethylation of cysteine residues as fixed modification and N-terminal acetylation and methionine oxidation as variable modifications), with the following exceptions: ‘Arg6;Lys4’ and ‘Arg10;Lys8’ were specified as medium and heavy modification labels, respectively; a maximum of three missed cleavages was allowed; ‘Phospho (STY)’, ‘Oxidation (M)’ and ‘Acetyl (Protein N-term)’ were selected as variable modifications; and the options ‘match between runs’ and ‘requantify’ were activated.

Ratios between labels (normalized by MaxQuant) were extracted from the Phospho(STY)Sites.txt file, reverse entries and potential contaminants were removed, and the list was filtered for at least two unique peptides per phosphosite. Phosphosite ratios were normalized to the respective ratio at the protein group level using the median ratio of all protein group ratios referenced in the ‘Protein group IDs’ column of the Phospho(STY)Sites.txt file. Resulting phospho site ratios were filtered so that only sites with valid values in >=3 of 4 replicates were retained. Significance of differential regulation was tested using linear models implemented in the LIMMA package and the resulting fold changes and significance level were plotted. Data analysis was performed in Python 3.10 using the autoprot wrapper ^94^ for data analysis and plotting. All original code for MS data analysis has been deposited at Zenodo (<https://doi.org/10.5281/zenodo.15350913>).
