## Supplementary material for "Microtubule anchoring and coupling of CD20 to the RhoA/Rock1 pathway": reagents and Tools

**Structured Methods - Reagents and Tools Table**

| **Cells** |  |  |
| --- | --- | --- |
| Healthy Donor B cells | Uniklinik Freiburg | buffy coats of Transfusion Medicine |
| Ramos Burkitt lymphoma | American Type Culture Collection | CRL-1923 |
| **Antibodies** |  |  |
| Anti CD20 mc 2H7 | Biolegend | 302314 |
| Anti CD19 mc HIB19 | Biolegend | 302216 |
| Anti CD32b mc S18005H | Biolegend | 398306 |
| Anti IgM mc MHM88 | Biolegend | 314536 |
| Anti IgD mc IA6-2 | Bolegend | 348222 |
| Anti CD83 mc HB15e | Biolegend | 305326 |
| Anti CD22 mc HIP22 | Biolegend | 302514 |
| Anti CD45 mc HI30 | Biolegend | 304016 |
| Anti CD69 mc FN50 | Biolegend | 310912 |
| Anti GEF-H1 mc 55B6 | Cell Signaling | 4076 |
| Anti Rock1 pc | proteintech | 21850-1-AP |
| Anti 14-3-3 pc | proteintech | 14881-1-AP |
| Anti pGEF-H1 S886 pc | invitrogen | PA5-105221 |
| Anti pRhoA S188 pc | BIOSS | Bs 5330R |
| Anti RhoA mc 67B9 | Cell Signaling | 2117 |
| Anti pRock1 Y914 pc | invitrogen | PA5-105054 |
| Anti CD27 mc M-T271 | Biolegend | 356412 |
| anti CXCR4 mc 12G5 | Biolegend | 306514 |
| Anti HLA1 mc W6/32 | Biolegend | 311402 |
| Anti β-actin mc 13E5 | Cell signaling | 4970 |
| Anti a-tubulin pc | Sigma Aldrich | T5168 |
| Anti mouse-A555 IgG mc | invitrogen | A32727 |
| Anti rabbit pc-HRP | Cell Signaling | 7076 |
| Anti mouse pc-HRP | Cell Signaling | 7074 |
| Anti PKCdelta pc | Cell Signaling | 2058 |
| Anti flag mc D6W5B | Cell Signaling | 14793 |
| Anti flag mc M2 | Sigma | F3165 |
| Phospho-PKCδ (Thr505) Antibody pc | Cell Signaling | 9374 |
| **Oligonucleotides and other sequence-based reagents** |  |  |
| Geneblock CD20 N-term | Intergrated DNA technologies (IDT) | 397bp, see ET1 |
| Geneblock CD20 C-term | IDT | 258bp, see ET1 |
| Geneblock CD20 wt | IDT | 482bp, see ET1 |
| CD20 KO Exon 3 | IDT | gccctattgctatgcaatctgg |
| pLenti:PKCdelta | Addgene | Plasmid #52961 |
| CD32b KO Exon 4 | IDT | agcacgttgatccactgggg |
| pGEX-4T1-RhoA G17A | Addgene | Plasmid #69357 |
| **Chemicals, Enzymes and other reagents** |  |  |
| Proximity ligation assay, probemaker, plus, minus | Sigma-Aldrich | Duo92009-1KT Duo92010-1KT |
| PLA detection in situ orange | Sigma Aldrich | Duo92007-100RXN |
| Fc blocker | Biolegend | 422302 |
| Glutathione Agarose | Thermo Fisher | 25237 |
| ECL Solution Femto SS | Thermo Fisher | 34096 |
| Rituximab | Hoffmann-La Roche AG | CH |
| docetaxel trihydrate  RP-56976 | MedChemExpress | 114977-28-5 |
| Nocodazole, R17934 | Selleck-Chemicals | 2775 |
| LipoJet™ In Vitro Transfection Kit | SignaGen | Cat #: SL100468 |
| **Software and Database** |  |  |
| Prism 10 | GraphPad | Dotmatics |
| CellProfiler 4.2.5 | CellProfiler | https://cellprofiler.org/ |
| Software ZEN Black | ZEISS | 2.3 SP1 FP3 |
| Inkscape 1.4.2 | Inkscape | https://inkscape.org/ |
| Snapgene 4.3.11 | Snapgene.com | Dotmatics |
| ImageStudioLite | LICOR | https://licorbio.com/image-studio-lite |
| FlowJo | Tristar | https://www.flowjo.com/ |
| MaxQuant version 2.6.7.0 | MaxQuant | MaxQuant.org |
| Limma package, | R | https://www.r-project.org/ |
| clusterProfiler package version 4.12.6 | R | https://www.r-project.org/ |
| edge R, version 4.4.0 | R | https://www.r-project.org/ |
| PRIDE data | ProteomXchange | https://www.proteomexchange.org/ |
| Molecular Signatures Database MSigDB, version 7 | GSEA | Broad Institute |
| GEO accession GSE300605 | NCBI | https://www.ncbi.nlm.nih.gov/geo/query/acc.cgi?acc=GSE300605) |
| **Other** |  |  |
| Airyscan Zeiss | ZEISS |  |
| DMi8 microscope | LEICA |  |
| ChromoTek DYKDDDDK Fab-Trap® Agarose | Proteintech | **Cat No.** ffa |
| Attune NxT Flow Cytometer | Thermo Fisher Scientific |  |
| UltiMate 3000 RSLCnano HPLC system | Thermo Fisher Scientific | Dreieich, Germany |
| QExactive Plus mass spectrometer  nanoEase M/Z Symmetry C18 Trap | Thermo Fisher Scientific | Bremen, Germany  Waters Corporation, Milford, MA |
| Q-page tgn percast gel (12 wells; 10%), smobio | 7-biosciences | QP5220 |
| IBIDI uncoated 8 well high glass bottom chambers | IBIDI | Cat. No 80807 |
| Nanospray Flex ion source with DirectJunction adaptor | Thermo Fisher Scientific |  |
| PTFE slides | Fisher Scientific | NC9811708 |
| Qubit 3.0; | life technologies |  |
| RNeasy plus kit | Qiagen | cat# 74143 |
| NEBNext® Ultra™ II Directional RNA Library Prep Kit | NEB | cat # E7760L |
| NextSeq® 500/550 High Output Kit v2 | Illumina | Cat. FC-420-1001-4 |
